# Mapping *trans*-eQTLs at single-cell resolution using Latent Interaction Variational Inference

**DOI:** 10.64898/2026.02.04.703363

**Authors:** Danai Vagiaki, Tobias Heinen, Manu Saraswat, Brian Clarke, Oliver Stegle

## Abstract

Single-cell expression quantitative trait loci (eQTL) studies hold promise for linking genetic variants to changes in gene expression in individual cells, thereby offering deeper insights into the genetic basis of human traits and diseases. Existing methods for mapping *cis* effects rely on predefined cell types, pseudobulk aggregation, or single-gene association tests, and are thus limited in their ability to detect complex *trans* effects that act across gene networks and cell-type subpopulations. Here, we present ***L****atent* ***I****nteraction* ***V****ariational* ***I****nference* (*LIVI*), an interpretable deep learning framework for efficient mapping of *trans* genetic effects at cellular resolution. LIVI employs a structured sparse variational autoencoder architecture to decompose observed gene expression profiles into cell-state- and donor-specific variation. The model enables efficient donor-level association testing while retaining single-cell resolution and interpretation. Applied to the OneK1K dataset, LIVI discovered a greater number of *trans*-eQTLs than alternative latent variable methods, yielded *trans*-eQTLs missed by conventional single-gene testing strategies and revealed the cell types and genes implicated in polygenic risk for autoimmune diseases. All in all, we demonstrate that LIVI is a powerful approach for linking genetics to gene regulation and through to human traits.

## Introduction

Expression quantitative trait loci (eQTL) studies, which probe for associations between genetic polymorphisms and variation in gene expression levels, can elucidate the molecular consequences of genetic variants implicated in human traits or disease. Indeed, eQTL studies in large cohorts have shown that trait-associated genetic variants discovered in genome-wide association studies (GWAS) can induce *cis*-effects on nearby genes and affect downstream cellular programs in *trans* [1, 2, 3, 4, 5]. Conventional eQTL mapping strategies employ linear association tests between the expression of individual genes and proximal, putatively *cis*-acting genetic variants. Advances in single-cell RNA sequencing (scRNA-seq) technologies and their application to population-scale cohorts have greatly increased the resolution of eQTL studies, enabling the discovery of eQTLs specific to different cell types, or even within cell-type subpopulations (*cell states*) [6, 7, 8, 9]. Proposed methods for single-cell resolution *cis*-eQTL mapping include linear mixed models [7, 10] that capture variation at the level of individual cells, thereby mitigating the need to define cell types *a priori*. Additionally, methods such as [11] have demonstrated advantages when mapping genetic effects, while appropriately modeling the count nature of the input data. However, these current methods do not scale well to datasets comprising millions of cells, as they are now becoming available through resources such as [8]. Most importantly, existing methods are not practical for mapping *trans* effects, which require performing individual association tests between all possible variant–gene pairs, imposing both a significant computational and a multiple testing burden. In addition, *trans*-eQTLs are notoriously more difficult to map than *cis*-eQTLs, due not only to the larger number of tests conducted, but also because the effect sizes tend to be smaller than those of *cis* effects [12].

Besides the computational and statistical cost, mapping *trans*-eQTLs using single-gene models does not reflect the underlying biology, particularly the complexity of cellular signaling cascades and gene regulatory networks, which imply that *trans* effects of a given variant are likely to manifest across multiple downstream genes. Indeed, strategies that test for associations between genetic variants and groups of genes can reduce the number of required tests by several orders of magnitude, and aid the detection of weak *trans* effects by aggregating them into stronger gene set-level signals [13]. To capture correlated genetic effects in bulk eQTL studies, prior work has considered latent factor or topic models, which learn latent groupings of co-regulated genes [14, 15]. Latent variable models (LVMs) also have a long tradition in single-cell genomics [16, 17, 18], and more recently have been adapted to model genetic effects on single-cell expression [19]. However, SURGE [19] does not model the count nature of the input data and adopts single-gene testing, thus ignoring gene regulatory dependencies. Moreover, genetic variants are already used during the inference of the latent variables, creating the need for a computationally expensive permutation scheme, in order to obtain calibrated test statistics. To avoid statistical circularity in downstream association testing with donor phenotypes, recent methods aimed to learn latent variables informed solely by donor identity in a generic fashion, without directly incorporating other donor characteristics [20, 21]. However, scITD [20] captures interindividual variation in scRNA-seq data at the level of discrete, predefined cell types rather than single cells, and MrVI [21] is not designed to identify small *trans*-eQTL effects, despite the fact that it models raw scRNA-seq counts (Table S1).

Here, we propose ***L****atent* ***I****nteraction* ***V****ariational* ***I****nference* (*LIVI*), a computational framework specifically designed for mapping *trans*-eQTLs in population-scale single-cell cohorts. LIVI combines the scalability of variational autoencoders (VAE) with the interpretability of linear latent factor models, allowing for efficient mapping of *trans* effects at single-cell resolution. The model builds on a VAE [22, 23] with a linear decoder [24] to jointly model interindividual and single-cell variation in gene expression data from multiple individuals, experiments and cell types (Fig.1). LIVI does not rely on predefined discrete cell type annotations and instead leverages the inferred latent components to derive the most relevant groupings of cells to discover genetic effects. Additionally, LIVI factors capture sets of co-regulated genes. Because of these properties, LIVI can flexibly aggregate and share information across both cells and genes (Fig. S1). Importantly, the model takes raw single-cell gene counts as input, precluding biases introduced by normalization and aggregation procedures.

We applied LIVI to the currently largest publicly available scRNA-seq dataset of genetically diverse human donors, comprising over one million peripheral blood mononuclear cells (PBMCs) from almost a thousand individuals [8]. The model identified *trans*-eQTLs mediated by disease-associated variants that were consistent with results from a well-powered bulk RNA-seq meta-analysis [25] and pinpointed activity of these eQTLs at single-cell resolution. Notably, LIVI outperforms other latent variable methods in detecting *trans* genetic effects and enables the discovery of *trans*-eQTLs missed by conventional gene-level testing approaches. Lastly, we used LIVI to identify cell types and genes implicated in the polygenic risk for autoimmune diseases, recovering many known disease-gene and disease-cell type associations.

## Results

### The LIVI model

LIVI is a probabilistic model for scRNA-seq data collected from a large population of individuals that explicitly models cell-state variation, donor-specific effects, as well as their interactions. At its core, LIVI builds on variational autoencoders (VAE) [22, 23], employing structured linear decoding networks that are inherently interpretable [24]. The resulting model has properties that resemble classical factor analysis, where the decoder is a factor loadings matrix (e.g. [26]).

For a single cell *i* from donor *y*, LIVI takes the donor identifier (ID) and the vector of observed expression counts for *M* genes, **x**_*i*_ ∈ ℤ^*M*^, as input, and decomposes the latter into canonical cell-state variation, cell-state-specific donor effects (*D* × *C*), and global donor effects on gene expression. Cell-state variation is encoded in a lower-dimensional cell-state latent space, 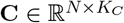, where *N* corresponds to the number of cells and *K*_*C*_ is the number of cell-state latent factors. *K*_*C*_ is typically small to encourage that **C** captures only canonical cell-state variability. Cell-state-specific donor effects are explained as a latent linear interaction between **C**, and a cell-state-specific donor embedding, 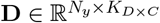, where *N*_*y*_ is the total number of individuals (donors), and *K*_*D*×*C*_ corresponds to the number of *D* × *C* latent factors, i.e., columns of the resulting interaction matrix (**CA**) ⊙ **D** (Fig. 1a). Here, ⊙ denotes the Hadamard (element-wise) product and 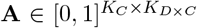 is a factor *assignment matrix* that “assigns” latent factors from 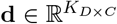 to cell-state factors, 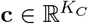. Intuitively, *D* factors capture donor effects, which are mapped onto the space of individual cells by the linear interaction term (**CA**) ⊙ **D**. A second donor embedding, 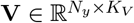, is included to account for donor latent variables causing broad changes in gene expression, such as population-structure. The latent spaces **C, D** × **C**, and **V** are decoded through separate linear decoders, enabling identification of gene programs associated specifically with cell states, cell-state–specific donor effects, and global donor effects. Moreover, to enhance the model’s ability to capture subtle cell-state-specific effects of individual SNPs, we use a larger number of *D* × *C* latent variables (*K*_*D*×*C*_ *» K*_*C*_ *> K*_*V*_) and decode those explicitly using a sparse linear decoder 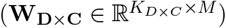 to portray gene regulatory programs. By contrast, global latent factors *V* that capture broad interindividual gene expression variation are decoded by a dense decoder 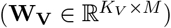 (Fig. 1a). Optionally, LIVI can account for donor or cell-level nuisance variables, such as age, sex and experimental batch (Methods). To increase power to detect *trans* effects, known *cis*-eQTLs may also be introduced to the model during training. By accounting for known *cis*-eQTLs, and modeling the effects of donor latent variables on all genes, regardless of genomic location, LIVI is inherently well suited to detecting *trans* genetic effects.

**Figure 1.**
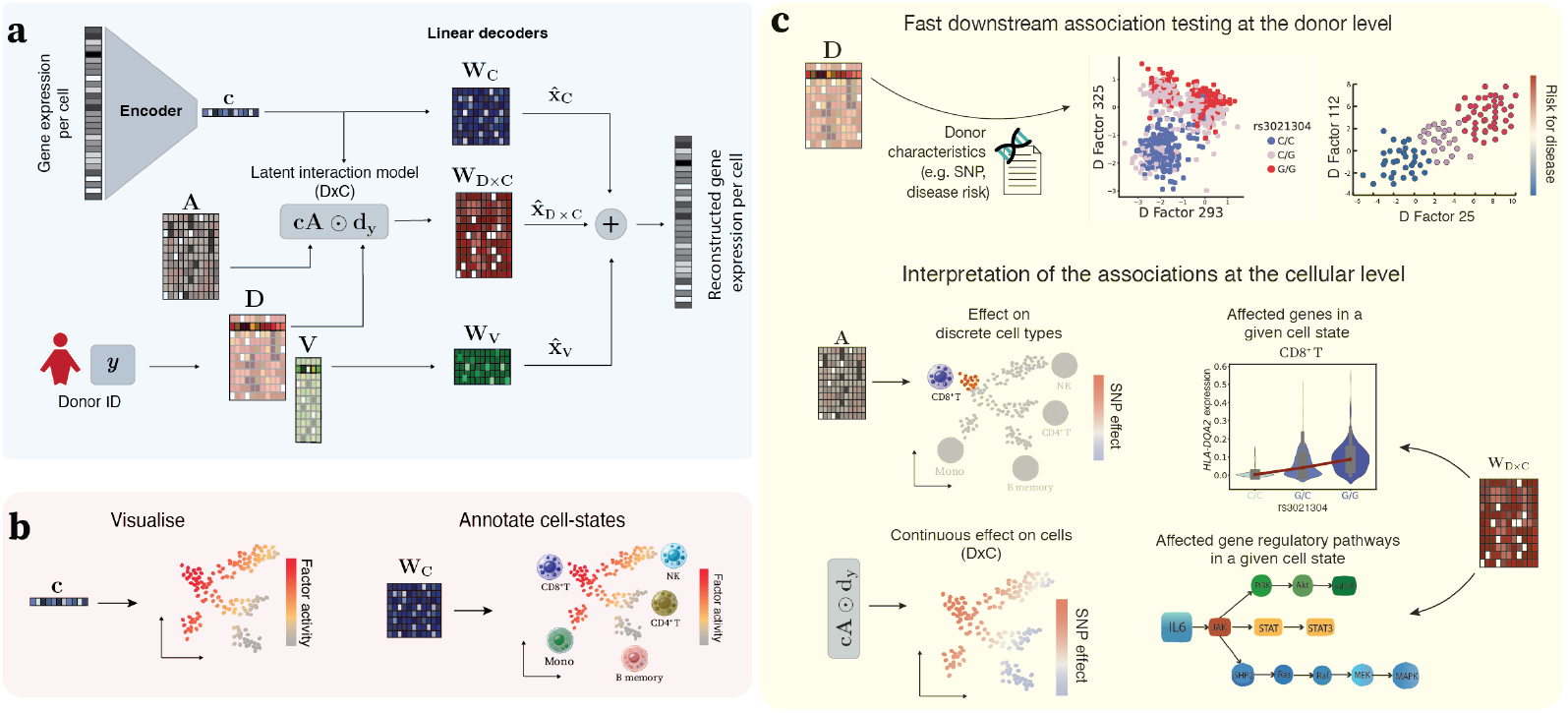
Overview of the LIVI model. **(a)** LIVI is an interpretable deep-learning framework for mapping *trans* effects of (disease-associated) genetic variants at the single-cell level. It decomposes the observed variation in gene expression into canonical cell states (**c**) and donor-specific effects (**D, V**). The donor embedding **D** captures cell-state-specific donor effects on gene expression, while **V** accounts for global sources of interindividual variation. Each of these latent representations is decoded by a dedicated linear decoder (with weights **W**_**C**_, **W**_**D**×**C**_, **W**_**V**_), which maps latent factors to individual genes. **(b)** LIVI’s cell-state factors can be used to identify and visualize cell states from scRNA-seq data. **(c)** After training, **D** factors can be used as phenotypes in a typical eQTL mapping framework (instead of individual genes) or probed for associations with other donor variables of interest. Discovered associations can be quantified and visualized either directly at the level of individual cells or in aggregate for a set of cells or cell types. Owing to the linear decoder architecture, associations with LIVI factors can be directly mapped to the implicated sets of genes, which can then be investigated for enrichment of specific cellular processes and gene regulatory pathways.

Once trained, LIVI enables a range of downstream association analyses (Fig. 1c). Because the donor factors are inferred without information on specific donor-level characteristics, such as SNP genotypes, they can be used as quantitative phenotypes to test for genetic effects without the risk of circularity (Methods). Following association testing at the donor level, the discovered effects can be projected back onto single cells via LIVI’s latent donor-cell-state interaction model (*D* × *C*), and the decoder weights can be inspected to identify the affected sets of genes (Methods). Besides single-variant association testing, donor factors *D* can be utilized to probe for cumulative effects of multiple variants on cellular programs, for example by testing polygenic risk scores (PRS). In principle, any donor-level phenotype of interest, such as disease status, can be assessed for association with LIVI’s donor factors, and subsequently mapped to affected cells and genes. Complementary to association testing, LIVI’s cell-state factors can serve as a dimensionality reduction and visualization tool for single-cell transcriptomics, similar to other latent variable models such as PCA or scVI [16] (Fig. 1b).

### Application of LIVI on a single-cell dataset of approximately one thousand individuals

#### LIVI decomposes observed gene-expression variation into interindividual and cell-state components

We applied LIVI to the OneK1K dataset [8], which comprises over one million peripheral blood mononuclear cells (PBMCs) from 981 donors with known cell type labels. Unless stated otherwise, we fit LIVI using 15 cell-state factors, 700 *D* factors and 5 *V* factors.

As anticipated, cell-state latent factors identified by LIVI explain cell type differences (Fig. 2a-b) to an extent that is comparable to conventional latent variable models that ignore the donor structure, such as PCA (Fig. S2). Reassuringly, the cell-state latent factors do not discriminate between donors, while the donor embeddings **D** and **V** do (Fig. S2), indicating that the model successfully disentangles latent variation attributable to either cell state or donor identity.

**Figure 2.**
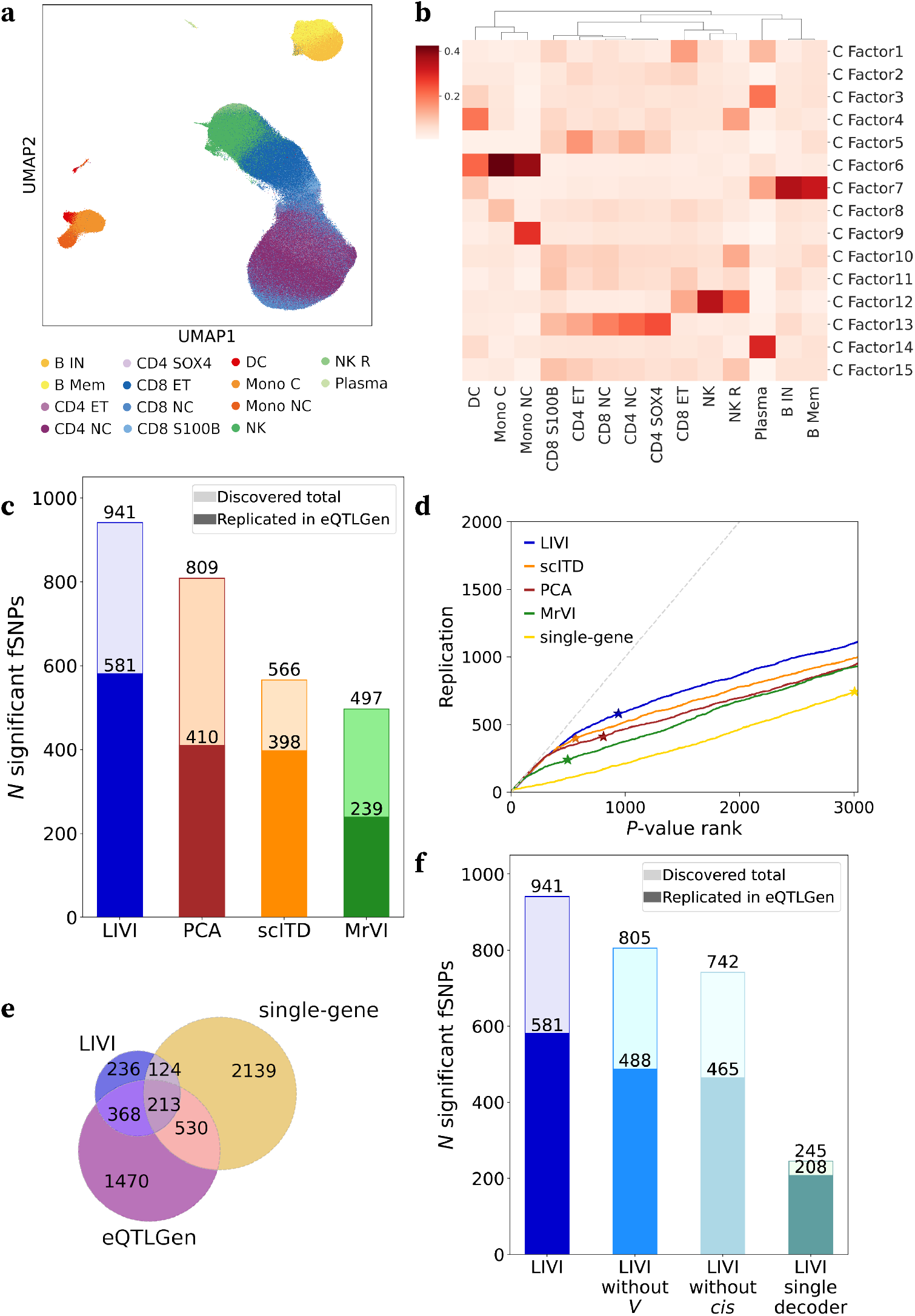
LIVI captures cell states, and outperforms alternative methods in *trans*-eQTL discovery. **(a)** UMAP representation of the cell-state latent space identified by LIVI. Color denotes the cell type labels from the original publication [8]. **(b)** Heatmap of cell-state factor values averaged across cells of different types, as annotated in the original publication. **(c)** Benchmark of alternative latent variable methods used for eQTL discovery. Shown is the number of variants in association with at least one latent variable estimated using LIVI, scITD [20], MrVI [21] or PCA (FDR < 5%). For each method, the number of latent variables yielding the largest number of discoveries was considered (700 for LIVI and scITD, 500 for MrVI and 100 for PCA; Fig. S11). Darker colors depict variants that were also identified (replicated) as significant *trans*-eQTLs in eQTLGen [25] (FDR < 5%). **(d)** SNP *p*-value rank (x-axis) vs. SNP replication count in eQTLGen [25] (y-axis) for associations obtained using latent variable methods as in **(c)**. In addition, we considered single-gene association testing for the 14,212 genes used to train LIVI. Stars indicate significance at FDR < 5%. **(e)** Overlap between SNPs discovered by LIVI vs. single-gene testing in the OneK1K [8] data and SNPs discovered in eQTLGen [25]. **(f)** Shown is the number of variants in association with at least one donor *D* factor for the full LIVI model (*LIVI*), as well as three reduced model formulations: one without global donor latent factors, **V** (*LIVI without V*), one without correction for known *cis*-eQTLs (*LIVI without cis*), and one with a single shared decoder for cell-state and donor factors (*LIVI single decoder*) (Methods). Darker colors depict variants replicated in eQTLGen [25].

A single cell-state factor may represent one cell type (e.g., factor 9 representing non-classical monocytes), or multiple related cell types (e.g. factor 13 activity across CD4^+^ T cells) (Fig. 2b). For datasets without *a priori* defined cell types, gene loadings for the cell-state factors may be utilized in a gene set enrichment analysis (GSEA) framework to annotate those factors based on known cell types from publicly available databases (Fig. S3; Methods).

The *D* × *C* factors estimated by LIVI represent gene programs that capture inter-individual variation modulated by cellular states. *D* × *C* factors are active in specific subsets of cells (Fig. S4), and the biological processes characterizing those cellular programs can be derived by GSEA on the loadings of the linear **W**_**D**×**C**_ decoder (Table S3; Figs. S5-S8). Lastly, we confirmed that the *D* × *C* factors are sparse, with a median of two genes (out of 14,212 genes) predominantly driving each factor (Fig. S9; Table S2; Methods).

### LIVI’s latent-space decomposition increases power to detect *trans*-eQTLs

We considered LIVI’s donor-aware latent space **D** as a set of quantitative traits in a classical eQTL analysis workflow [27, 28]. Briefly, we used a linear mixed model (Methods) to test each of the 700 latent *D* factors for associations with 9,305 GWAS-associated SNPs that were previously assessed for *trans*-eQTL effects in eQTLGen [25], a bulk blood RNA-seq eQTL meta-analysis study of 31,684 individuals. Even though eQTLGen is not a single-cell study, it is expected that genuine eQTL signals, even cell-type-specific effects, are detectable in a dataset of this size (32 times larger than the OneK1K dataset [8]).

In total, we identified 4,013 SNP-factor associations (FDR < 5%; Table S4), which are hereafter referred to as *fQTLs* (factor-QTLs) with corresponding *fSNPs* (factor-associated SNPs). These associations involved 941 unique fSNPs (Fig. 2c) and 323 unique donor *D* factors. Most fSNPs were associated with a single *D* factor and vice versa (Fig. S10). The activity of those *D* factors in different cell states can be assessed by the assignment matrix **A** (Fig. 1a). This analysis revealed that most of the fQTLs identified by LIVI are cell-state-specific (Fig. S10).

For comparison, we applied the identical genetic analysis workflow to quantitative traits derived from alternative latent variable models that, like LIVI, do not use donor genotypes during training, including principal component analysis (PCA), MrVI [21], and scITD [20] (Figs. 2c-d and S11). LIVI identified the largest number of variants with at least one fQTL (Figs. 2c and S11). Additionally, 61.7% of fQTL variants discovered by LIVI replicated in eQTLGen, whereas the number of replicated variants was lower for alternative methods (Fig. 2c-d). We also note that while the number of discoveries depends on the number of latent variables for all methods considered, LIVI consistently recovered more associations across a range of latent variable settings (Fig. S11).

Although conceptually distinct and thus not directly comparable, we also evaluated the aforementioned latent variable methods against a single-gene mapping that is conventionally applied in eQTL studies, such as eQTLGen. This conventional approach yielded a larger number of eQTLs (3,006 eSNPs at FDR < 5%; Table S5) than any of the latent variable models; however, fewer of these replicated in eQTLGen (Fig. 2e). Notably, the difference in the number of discoveries disappeared when applying an alternative multiple-testing correction that accounts for the tremendous difference in the number of traits and therefore tests conducted by single-gene association testing (198,968 genes x cell types) vs. latent factor testing strategies such as LIVI (700 latent variables) (Fig. S11; Methods).

To assess the robustness of LIVI, we compared significant associations across multiple random initializations, finding that results were consistent across runs (Fig. S12; Methods). We also explored how LIVI behaves across different dataset sizes. Smaller datasets yielded no significant associations, suggesting that the model does not overfit, while larger datasets yielded a growing number of discoveries, without, however, reaching saturation at the size of OneK1K (Fig. S13).

Lastly, we examined the impact of the various model components on the power to detect fSNPs by training three reduced model formulations: first, a model that decodes cell-state and donor factors jointly; second, a model without global donor embedding, **V**; and finally, a model without correction for known *cis*-eQTLs (Methods). Our experiments show that the full LIVI model recovers a larger number of SNP-factor associations than any of the reduced models. Importantly, separately decoding cell-state-driven and donor-driven latent sources of variation is key to detecting SNP effects. (Figs. 2f and S14).

### LIVI discovers *trans*-eQTLs missed by conventional single-gene testing approaches

To further explore the advantage of LIVI for eQTL discovery, we considered the 604 variants found to be associated with LIVI factors (fSNPs), but not discovered when conducting single-gene tests in discrete cell types. We explore here two hypothesized scenarios in which LIVI provides a power advantage, namely for associations that either (i) act in cell-state space in a manner that does not align with discrete cell types, and/or (ii) involve multiple genes.

Indeed, we estimated that 47.2% of the 604 variants discovered exclusively by LIVI act in continuous cell states (Fig. S15; Methods). An example of such an fSNP is rs1610677, located in the human leukocyte antigen (*HLA*) locus and known to be associated with rheumatoid arthritis (RA) and hypothyroidism [29]. rs1610677 is a known *cis*-eQTL for *HLA-A* and *ZFP57* in whole blood [25, 30]. rs1610677 was also reported as a *trans*-eQTL in eQTLGen (FDR < 5%) [25]; however, these associations were not replicated by our single-gene testing analysis in the OneK1K dataset (FDR < 5%; Table S5, Fig. S16). Using LIVI, we identified an association between the rs1610677 T allele and D × C_287_, which is primarily driven by *KLRB1* expression in CD4^+^ T cells differentiating from naive (CD4 NC) to effector memory (CD4 ET) phenotype (Fig. 3a-b). Along this trajectory, we defined the cell population with high D × C_287_ activity 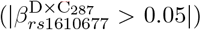 as the *LIVI subset*. Single-gene tests in CD4 NC, CD4 ET or the *LIVI subset* did not detect an association between rs1610677 and *KLRB1* expression at 5% FDR. Nevertheless, nominal evidence for association (unadjusted *p* < 0.05) was observed in CD4 NC (Table S6) and in the *LIVI subset*, with the strongest signal in the *LIVI subset* (Fig. 3b).

**Figure 3.**
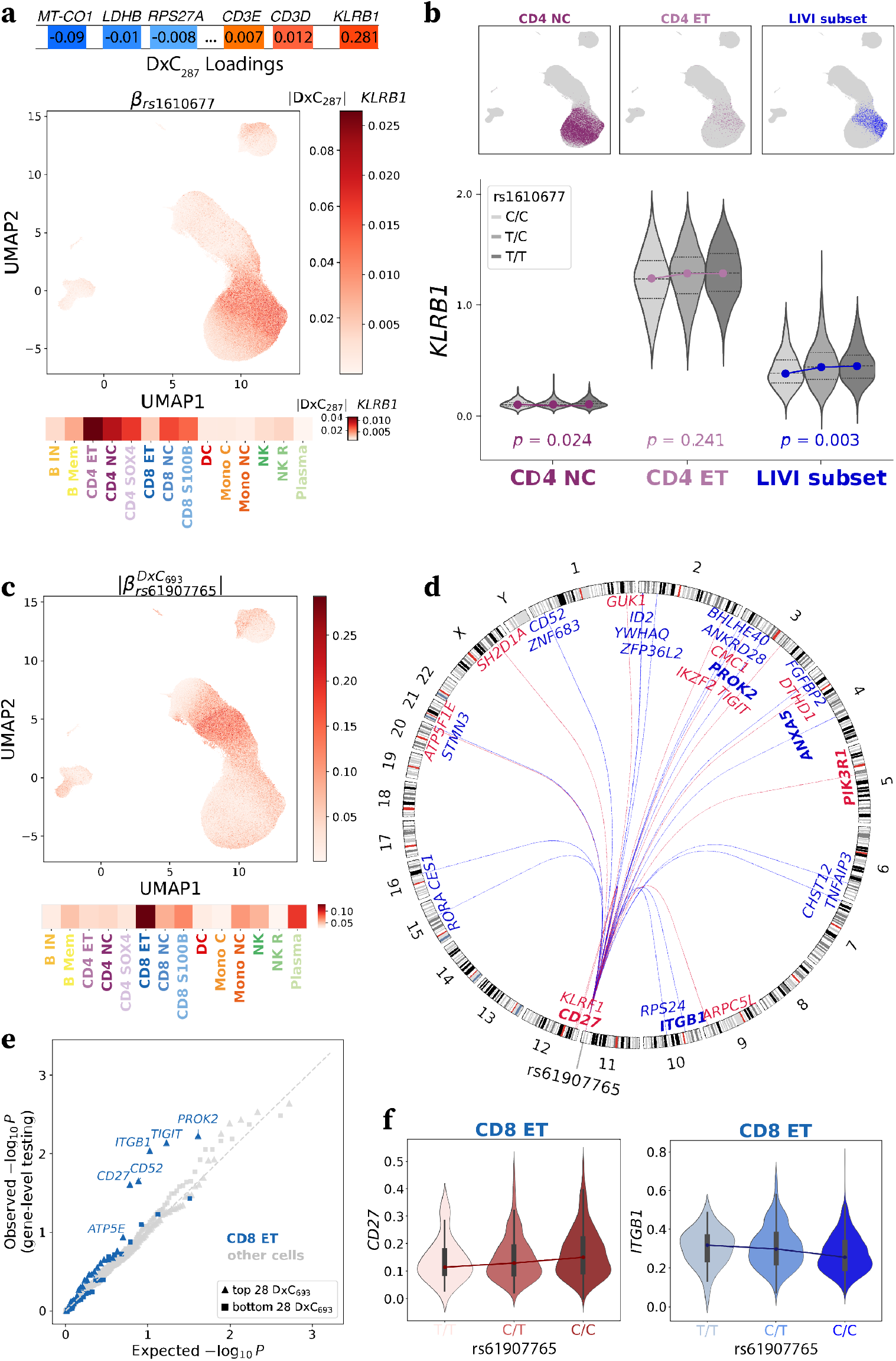
LIVI discovers *trans* acting variants missed by single-gene testing. **(a)** Top: Gene loadings for D × C_287_. Middle: UMAP of LIVI’s cell-state latent factors. Color denotes the estimated absolute effect of rs1610677 on D × C_287_, and specifically the estimated effect on the top-loaded gene, *KLRB1*, in each cell. Bottom: Estimated absolute effect of rs1610677 on D C_287_ and estimated effect on *KLRB1* averaged across cells of different types. **(b)** Top: UMAP of LIVI’s cell-state latent factors. Color denotes cells annotated as CD4 NC (left; dark purple) or CD4 ET (middle; light purple) in the original publication [8], or cells distinguished by high activity of D × C_287_ (*LVI subset*; right; blue). Bottom: Distribution of the observed log-normalized expression of *KLRB1*, pseudobulked across CD4 NC, CD4 ET or *LVI subset* cells for each donor and stratified by donor genotype at rs1610677. In each violin plot black solid lines indicate the median, while dotted lines indicate the interquartile range (IQR). Single-gene test association *p*-values between rs1610677 and *KLRB1* in each cell population are noted. **(c)** Top: UMAP of LIVI’s cell-state latent factors. Color denotes the estimated absolute effect of rs61907765 at the single-cell level. Bottom: Estimated absolute effect of rs61907765 on D × C_693_ averaged across cells of different types. **(d)** Circos plot illustrating the genomic locations of the top 28 genes for D × C_693_, which is associated with rs61907765 (a SNP located on chromosome 11). Gene names and connected lines are colored according to the estimated effect of the SNP on the expression of the respective gene: red denotes positive effect; blue negative effect. Genes belonging to GO:0043066 (*Negative Regulation Of Apoptotic Process*) are marked in bold. **(e)** Q-Qplot of expected vs. observed −*log*_10_-transformed *p*-value for association tests between rs61907765 and the 28 genes with the highest (triangles), as well as the lowest (squares) absolute loadings for D × C_693_, evaluated using single-gene tests in different cell types. Association *p*-values in CD8 ET cells marked in blue; association tests in other cell types shown in grey. **(f)** Distribution of the observed log-normalized expression of *CD27* (left) and *ITGB1* (right), pseudobulked across CD8 ET cells for each donor and stratified by donor genotype at rs61907765. Black boxes indicate the interquartile range (IQR), while whiskers mark the 1.5×IQR threshold.

*KLRB1* expression is one of the early markers of naive CD4^+^ T cells transitioning to memory CD4^+^ T cells [31]. We speculate that an increase in *HLA-A* expression, observed in the presence of the T allele of rs1610677 [30], could enhance T-cell receptor (TCR) signaling and eventually CD4^+^ T cell activation, leading to increased *KLRB1* expression. Consistent with this, *CD3D* and *CD3E*, encoding for the *δ* and *ϵ* TCR subunits, are among the top-loaded genes on the same *D* × *C* factor (Figs. 3a and S16). Alternatively, an epigenetic route for the inferred *trans*-eQTL is also plausible: the T allele of rs1610677 has a negative effect on the expression of *ZFP57* [30], a member of the KZFP family (zinc finger protein containing a KRAB domain) [30], which engages in the establishment and maintenance of repressive chromatin marks, such as H3K9me3. In particular, ZFP57 maintains methylation by binding to methylated 5’-TGCCGC-3’ consensus sequences. MEME/FIMO [32] analysis (Methods) identified two such motifs in the *KLRB1* promoter at positions chr12:9750774-9750780 and chr12:9761146-9761152 (GRCh37, reverse strand). Thus, ZFP57 could be involved in the maintenance of *KLRB1* promoter methylation status, implying that a decrease in *ZFP57* expression could lead to *KLRB1* hypomethylation, and thus increased expression, especially if permissive histone marks are also present. In fact, a link between decreased methylation and increased mRNA levels of *KLRB1* in tumor tissues has already been proposed [33].

To investigate our second hypothesis, that LIVI is better suited to discover SNP effects on multiple genes, we compared the number of associated eGenes in eQTLGen for fSNPs discovered by LIVI in OneK1K vs. eSNPs discovered by single-gene testing in OneK1K. This comparison confirmed that fSNPs affect a larger number of genes on average (Fig. S15). An example of such an fSNP is rs61907765, a variant known to be associated with celiac disease [34, 35] and psoriasis [36]. rs61907765 falls within the enhancer element ENSR11 D8D6G and is a known *cis*-eQTL for *ETS1* [25], a transcription factor gene known to be involved in the differentiation, survival and proliferation of lymphoid cells. In our analysis, rs61907765 was estimated to have an effect in effector memory CD8^+^ T cells (CD8 ET) (Fig. 3c) on the expression of 28 genes explaining D × C_693_ (Fig. 3d; Table S2). Three of those genes, including the top loaded gene, *ZNF683*, were also found to be associated with rs61907765 in eQTLGen [25] with the same direction of effect size (Fig. S17). Besides those three genes, D × C_693_ genes were enriched for *Negative Regulation Of Apoptotic Process* (GO:0043066) (*ITGB1, ANXA5, PROK2, CD27, PIK3R1*; *p* = 7 × 10^−4^; Table S3). LIVI identified a negative association between the C allele of rs61907765 and the expression of *ITGB1, ANXA5, PROK2*, as well as a positive association between the same allele and the expression of *CD27* and *PIK3R1* in effector memory CD8^+^ T cells (CD8 ET) (Figs. 3c-d and S18).

To understand why those associations were missed by single-gene testing, we inspected the single-gene association results with rs61907765 (Table S7). Reassuringly, the evidence for association between rs61907765 and the top genes of D × C_693_ was stronger in CD8 ET cells than in other cell types. This enrichment was confined to the top D × C_693_ genes and did not extend to genes with small loadings on the same factor (Fig. 3e). These observations suggest that LIVI correctly estimated those specific genes as affected by rs61907765 in CD8 ET cells. Notably, associations between rs61907765 and three genes belonging to the *Negative Regulation Of Apoptotic Process* (GO:0043066) pathway were nominally significant (uncorrected *p*-values: *PROK2*: *p* = 0.0059; *ITGB1*: *p* = 0.0092; *CD27*: *p* = 0.025) (Fig. 3e; Table S7). Taken together, rs61907765 likely influences the expression of *PROK2, ITGB1* and *CD27*, but the increased multiple-testing burden inherent to single-gene analyses obscured these associations. By reducing the number of tests and pooling weaker single-gene effects into effects on gene programs defined by *D* × *C* factors, LIVI is able to recover those associations.

### LIVI uncovers gene-program-level effects of known eSNPs

Besides uncovering previously missed *trans*-acting variants, LIVI can also furnish complementary evidence for variants detectable via single-gene tests. Associations discovered by single-gene tests can be difficult to interpret, especially when a genetic variant is found to be associated with only one gene, as this precludes the identification of the full gene regulatory pathways implicated. Since LIVI is tailored to detect effects on groups of genes, we investigated whether it could enhance the interpretation of such associations. In total, 337 SNPs were detected by both gene-level and factor-level tests; 45% of these (152) were associated with only one gene in single-gene analyses. Among these 152 single-gene-associated variants, we focused on the 70 located outside chromosome six, thereby excluding the *HLA* locus, as their consequences in immune cells are more challenging to interpret.

One of those variants is rs12550612, a SNP on chromosome eight known to be associated with blood traits, such as leukocyte quantity, myeloid leukocyte count [37] and neutrophil count [38, 39]. Single-gene association testing in the OneK1K dataset identified a negative association between the G allele of rs12550612 and the expression of *SLMO2* /*PRELID3B* in CD4 NC cells (*β* = −0.13; *p* = 3.06 × 10^−7^; Table S5), whereas no *trans* effects for rs12550612 were observed in eQTLGen [25]. Association testing with LIVI linked rs12550612 to the expression of 17 genes in CD4 NC (Fig. 4a-b). The associated LIVI factor, D × C_232_, distinguishes a cell population within CD4 NC 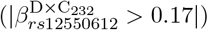, which we again refer to as *LIVI subset*. The 17 genes driving D × C_232_ did not include *SLMO2* (Fig. S19; Table S2). To investigate the discrepancy in the mapped genes between conventional single-gene vs. LIVI testing, we compared the association *p*-values obtained from single-gene tests for rs12550612 in the *LIVI subset* with those from the entire CD4 NC population (Fig. 4c). In the *LIVI subset*, rs12550612 was significantly associated with the expression of *SOCS3* (*p* = 7.8 × 10^−7^; FDR < 5%), but not *SLMO2* (*p* = 0.3; FDR < 5%) (Fig. 4c). In fact, *SOCS3* has the largest absolute loading on D × C_232_ (Fig. S19). Moreover, other genes explaining D × C_232_, such as *MYC, SOCS1, PIM1*, and *PIM3*, were nominally significant (unadjusted *p*< 0.05) in both the *LIVI subset* and the full CD4 NC population (Fig. 4c; Table S8). These findings demonstrate that LIVI can resolve distinct *trans*-eQTL effects in cell states that are more granular than canonical cell types.

**Figure 4.**
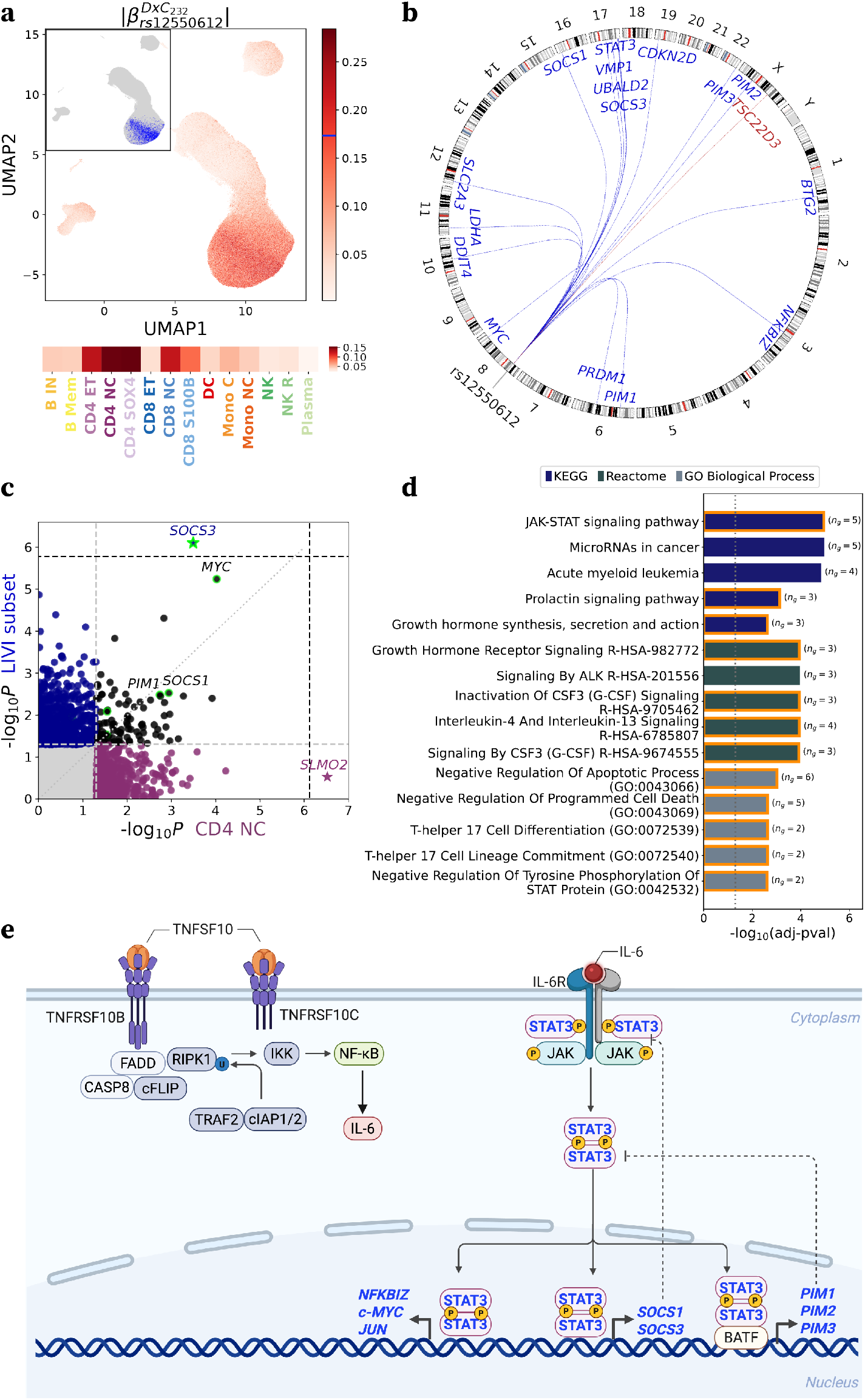
LIVI uncovers gene-program-level effects of known eSNPs. **(a)** Top: UMAP of LIVI’s cell-state latent factors, displaying the estimated absolute effect of rs12550612 on D × C_232_ at the single-cell level. The blue line on the colorbar denotes the D × C_232_ activity cut-off for cells defined as *LIVI subset*, which are also highlighed in blue in the inset. Bottom: Estimated absolute effect of rs12550612 on D × C_232_ averaged across cells of different types from the original publication [8]. **(b)** Circos plot illustrating the genomic locations of the top 17 genes for D × C_232_, which is associated with rs12550612, located on chromosome 8. Gene names and corresponding connecting lines are colored according to the estimated direction of effect (red: positive, blue: negative) of the SNP on the expression of the respective gene. **(c)** Scatterplot of −*log*_10_-transformed *p*-value of association tests between rs12550612 and individual genes in CD4 NC cells (x-axis) vs. *LIVI subset* (y-axis). The dotted grey line marks the diagonal. The dashed grey lines indicate the nominal significance threshold (*α* = 0.05). The dashed black lines indicate the significance threshold after FDR correction (FDR < 5%). Nominally significant SNP-gene associations are denoted with color (purple: CD4 NC, blue: *LIVI subset*, black: both). Stars indicate significant discoveries after multiple testing correction (FDR < 5%). Genes with high absolute loadings for D × C_232_ are highlighted in green. **(d)** Enrichment analysis results for the top 17 genes explaining D × C_232_. Each bar represents an enriched term, displayed along the y-axis, with the corresponding −*log*_10_(adjusted *p*-value) shown on the x-axis. The number of D × C_232_ top genes involved in each term is noted in parenthesis. Bar colors denote the source database used for enrichment analysis: KEGG, Reactome, Gene Ontology (GO) Biological Process. Bars corresponding to terms involving *SOCS3*, the top gene for D × C_232_, are framed in orange. **(e)** Illustrative diagram showing how the known *cis*-eQTL effect of rs12550612 on *TNFRSF10B*, a gene involved in the NF-*κB* pathway, can be connected to the *trans* effects identified by LIVI on genes implicated in JAK-STAT signaling. Briefly, NF-*κ*B activation can induce the expression of *IL6*. IL-6 acts as a ligand that activates the JAK-STAT signaling cascade, which culminates in the expression of various genes, including *MYC, NFKBIZ, SOCS1, SOCS3, PIM1, PIM2* and *PIM3*. JAK-STAT pathway implicated genes with high loadings for D × C_232_ are emphasized with bold, blue letters.

Next, we sought to investigate the *trans*-eQTL effect of rs12550612 on *SOCS3* by leveraging LIVI’s ability to identify groups of co-regulated genes. GSEA revealed that the 17 genes explaining D × C_232_ were most significantly enriched for genes involved in *JAK-STAT signaling pathway* and *MicroRNAs in cancer*; however only the former term included *SOCS3* (Fig. 4d). JAK-STAT signaling, specifically the JAK1-STAT3 signaling cascade, is known to be activated by the binding of interleukin-6 (IL-6) to its receptor [40]. Activated STAT translocates to the nucleus, where it regulates the expression of downstream target genes, such as *SOCS3, SOCS1, MYC, PIM1, PIM2*, and *PIM3* (Fig. 4e) – all of which had high loadings for D × C_232_. IL-6 production can be induced by the activation of nuclear factor kappa B (NF-*κ*B) through tumor necrosis factor–related apoptosis-inducing ligand - death receptor 5 (TRAIL-DR5) signaling [41, 42] (Fig. 4e). The A allele of rs12550612 is known to have a negative effect on the expression of *TNFRSF10B* [25, 30], the gene encoding for DR5. A reduction in the expression levels of *TNFRSF10B* could interfere with NF-*κ*B activation and IL-6 production. Consequently, IL-6 mediated activation of JAK-STAT signaling may be impaired, leading to decreased transcription of *SOCS3* and other STAT target genes. Indeed, LIVI estimates that the A allele of rs12550612 decreases the expression of the aforementioned genes (Fig. S19), which is in accordance with the observed eQTL effect using pseudobulk expression aggregation (Fig. S19).

### LIVI facilitates the identification of cell types and genes implicated in the polygenic risk for autoimmune diseases

In addition to *trans*-eQTL mapping of individual variants, LIVI *D* factors can be probed for associations with other genetic or non-genetic donor characteristics, including disease risk. To demonstrate this in the absence of phenotypic information for OneK1K, we used the PGS Catalog [43] to calculate polygenic risk scores (PGS/PRS) for seven autoimmune diseases and twelve blood traits (Methods). We also included height, hair color and myopia as negative controls traits (Methods), as these are not expected to be mediated by blood cell types. Association analysis with LIVI *D* factors resulted in 66 associations, covering six autoimmune diseases and three blood traits (FDR < 5%; Fig. 5a; Table S9). Reassuringly, there was no association between LIVI factors and the three negative control traits. In what follows, we focus on the six autoimmune diseases associated to LIVI factors: rheumatoid arthritis (RA), multiple sclerosis (MS), type 1 diabetes mellitus (T1D), psoriasis, ulcerative colitis (UC), and celiac disease (CeD).

**Figure 5.**
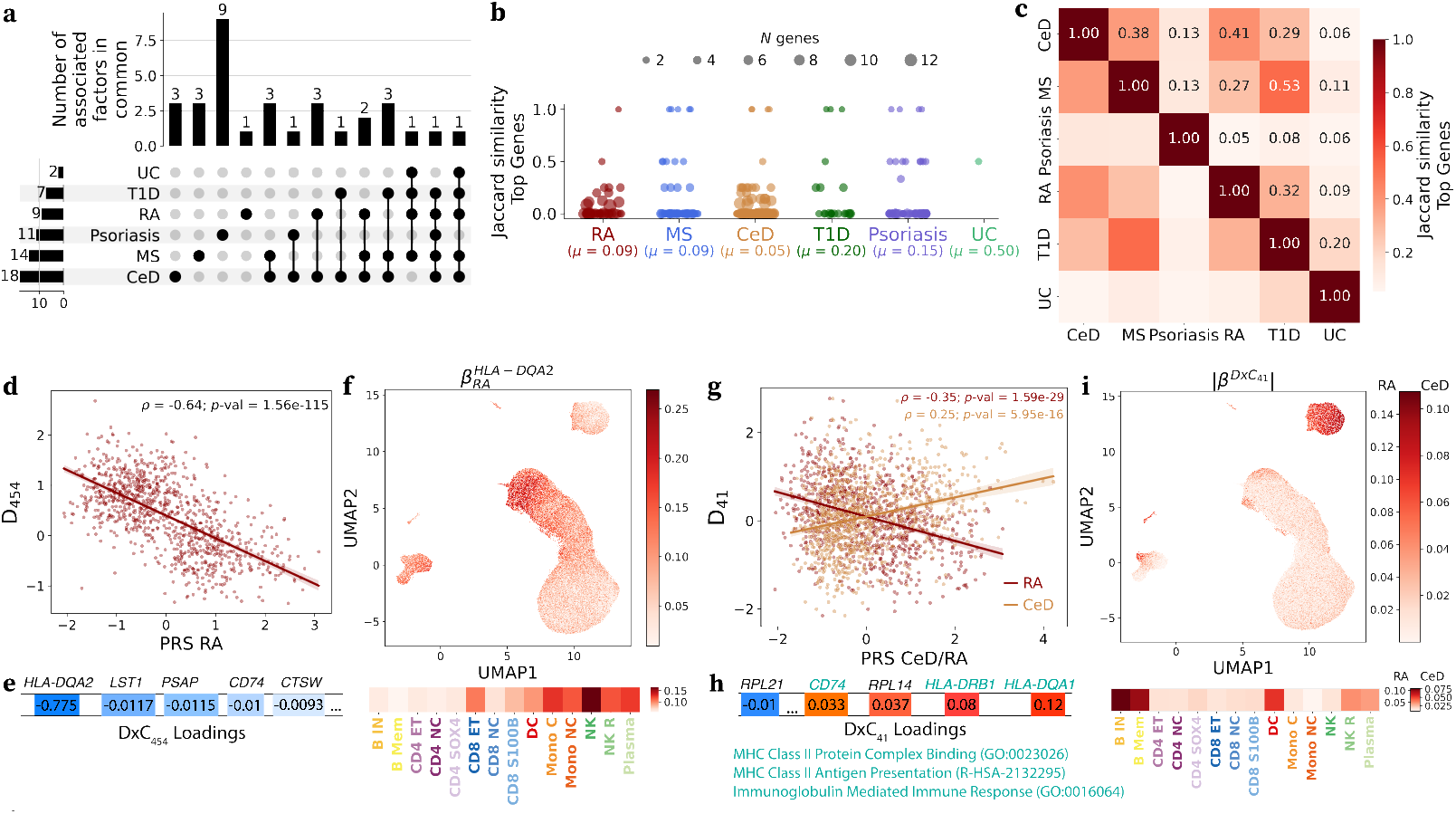
LIVI identifies cell states and genes involved in polygenic disease risk. **(a)** Upset plot of *D* factors associated the PRSs for autoimmune disease. **(b)** Jaccard similarities for pairs of factors associated with a given PRS, calculated based on the top loaded genes for those factors (Methods). The size of the dots corresponds to the number of genes considered for calculating the Jaccard similarity between two factors. **(c)** Jaccard similarities between PRSs, calculated based on the top loaded genes for all factors associated with a given PRS. **(d)** Scatterplot between PRS for RA vs. D_454_ activity in each donor. Spearman’s *ρ* between PRS for RA and D_454_ and corresponding *p*-value are noted. **(e)** Gene loadings for D × C_454_, which corresponds to D_454_ projected to the single-cell level. **(f)** Top: UMAP of LIVI’s cell-state latent factors. Color denotes the estimated effect of PRS for RA on *HLA-DQA2* at the single-cell level. Bottom: Estimated effect of PRS for RA on *HLA-DQA2* averaged across cells of different types from the original publication [8]. **(g)** Scatterplot between PRS for RA (bordeaux) or CeD (gold) and D_41_ values for each donor. Spearman’s *ρ* between PRSs and D_41_ and corresponding *p*-values are noted. **(h)** Gene loadings for D × C_41_, which corresponds to D_41_ projected to the single-cell level. Three genes involved in *MHC Class II Protein Complex Binding* (GO:0023026), *MHC Class II Antigen Presentation* (R-HSA-2132295), and *Immunoglobulin mediated Immune Response* (GO:0016064) are noted in turquoise. **(i)** Top: UMAP of LIVI’s cell-state latent factors. Color denotes the estimated absolute effect of PRS for RA or CeD at the single-cell level. Bottom: Estimated absolute effect of PRS for RA or CeD averaged across cells of different types from the original publication.

Similar to SNP-factor associations, most *D* factors were uniquely associated with a single PRS (Figs. 5a and S20). However, unlike SNP-factor associations, PRSs for autoimmune diseases tended to be associated with multiple factors (Figs. 5a and S20), likely reflecting that a PRS summarizes different genetic components of a given trait. Indeed, the associated *D* × *C* factors largely capture distinct gene programs (Fig. 5b). Likewise, factors associated with PRSs for different diseases were principally driven by distinct gene sets, except for MS and T1D, diseases that have been reported to co-occur in children and adolescents [44] (Fig. 5c). The few factors associated with PRS for multiple autoimmune diseases are explained by *HLA* genes, especially those coding for MHC class II proteins (Table S10). Of note, *HLA-DRB5* was shared between all six autoimmune diseases and *HLA-DRB1* among five PRSs: RA, T1D, MS, CeD, and Psoriasis (Table S10). *HLA-DRB1* alleles have been linked to the risk of RA, MS, T1D, and Psoriasis in previous studies [45, 46, 47, 48, 49]. Those relations could also explain the discovered association between *HLA-DRB5* and the same diseases, since *HLA-DRB5* and *HLA-DRB1* are in strong linkage disequilibrium (LD), and therefore typically inherited together [50].

The strongest and most statistically significant association was between D_454_ and the PRS for RA (*β* = −0.45; *p* < 1.23 × 10^−110^; Table S10, Spearman’s *ρ* = −0.64; *p* < 1.56 × 10^−115^; Fig. 5d). The corresponding factor D × C_454_ is primarily active in NK cells and explained by *HLA-DQA2* (Figs. 5d-f and S21). To the best of our knowledge, there is no known link between *HLA-DQA2* and RA predisposition. However, an accumulation of NK cells has been observed in inflamed synovial joints of RA patients [51, 52]. In addition, cell-type-specific association testing between PRS and individual genes replicated this strong association between *HLA-DQA2* and RA polygenic risk in NK cells (*β* = 1.22; *p* < 5.67 × 10^−48^; Table S11). Notably, PRSs for T1D and MS were also associated with D_454_ (Table S10; Fig. S21), and consequently *HLA-DQA2* in NK cells, and those associations were also detectable by single-gene testing (Table S11). Studies suggest that NK cells play a protective role in MS by inhibiting autoreactive T cells with pathogenic potential [53], and NK/CD4^+^ T cell ratio may serve as a prognostic indicator for MS disease activity [54]. In addition, reduced abundance and cytotoxicity of NK cells have been reported in T1D patients [55, 56, 57].

Besides strong associations between multiple PRSs and factors dominated by *HLA* genes – such as the one described above – we also obtained weaker, more disease-specific associations. For instance, PRSs for RA and CeD were both associated with D_41_ (Figs. 5g and S21). The corresponding D × C_41_ is explained by genes involved in *MHC Class II Antigen Presentation* (R-HSA-2132295) and *Immunoglobulin Mediated Immune Response* (GO:0016064), and is active in B cells (Fig. 5h-i). This supports the known pathophysiology of these diseases, which are driven by autoantibody production, such as anti-citrullinated protein antibodies (ACPA) and rheumatoid factor (RF) in the case of RA [58, 59] or anti–tissue transglutaminase (anti-tGT) in the case of CeD [60]. Interestingly, D_41_ was not associated with diseases in which the primary cause is not autoantibody production by B cell-derived plasma cells, such as MS [61, 62], T1D [63] or psoriasis [64].

## Discussion

Here, we have proposed *LIVI*, an interpretable latent variable model that enables scalable mapping of genetic effects at cellular resolution. LIVI builds on the VAE model [22], combined with a linear decoder architecture to encourage an interpretable mapping of latent to observed variables (genes), similar to the concept of factor analysis. LIVI directly models gene expression counts from single cells, and decomposes the observed gene expression into canonical cell-state variation, cell-state-specific donor effects and global donor effects, while optionally controlling for technical (batch) effects and known *cis*-eQTL effects.

In our benchmarks, we showed that LIVI outperforms alternative single-cell latent variable methods [20, 21] in identifying genetic effects. We also demonstrated its ability to detect genetic signals that involve multiple genes or that do not conform to traditional discrete cell type annotations, and are therefore missed by conventional single-gene testing approaches. Furthermore, we illustrated that LIVI enhances the interpretation of eSNPs discovered by single-gene tests by capturing pathway-level information. Additionally, LIVI can be utilized to perform association testing directly with disease risk, thereby broadening its applicability in understanding the molecular mechanisms underlying disease.

Contrary to alternative single-cell latent variable models designed to capture genetic effects [19], LIVI does not use the donor genotypes during model training, mitigating the risk of statistical circularity in downstream genotype-phenotype association testing. While conceptually related to other single-cell latent variable methods that infer a generic donor representation [20, 21], LIVI is unique in that it infers distinct gene regulatory programs to explain cell state vs. donor-specific variation, leading to increased power to detect genetic effects, as illustrated in our benchmarks. Similar to MrVI, LIVI also directly models single-cell count profiles as inputs.

The primary limitation of our method is that statistical significance is assessed at the factor level, and *p*-values cannot be assigned directly to associations of individual genes. However, our results demonstrate that analyzing the top-loaded genes within a factor can provide gene-level insights that are consistent with those gained through single-gene testing. Another non-trivial issue is choosing the right set of model hyperparameters, chief among these being the number of *D* factors. In typical machine learning tasks, e.g., supervised learning, training/validation loss serves as a suitable objective for hyperparameter optimization. However, eQTL studies commonly optimize for the number of discoveries [65], which is not directly evaluated during LIVI’s training. Here, we optimized key model parameters, including the number of *D* factors, for the OneK1K dataset. As the number of available population-scale scRNA-seq datasets increases [9, 66], application of LIVI to these will help establish guidelines for selecting optimal model hyperparameters based on the number of donors, expressed genes, and cell types.

A natural extension of LIVI in the era of multi-omics datasets would be to inform the cell-state latent space by additional omics modalities for the same cell, such as single-cell assay for transposase-accessible chromatin using sequencing (scATAC-seq) data. Besides offering a more holistic cell-state representation, this could also reveal relevant accessible genomic regions for each cell-state, which could in turn help to prioritize genetic variants for association testing: variants located inside cell-state-specific accessible regions can be assessed for effects on genes characterizing the same cell-state. Another area of interest is testing LIVI’s donor factors for associations with specific disease diagnoses. It is expected that the presence of the disease phenotype will have a stronger impact on transcriptomic signatures than genetic susceptibility alone. In this work, we provided evidence that LIVI factors capture genetic susceptibility to disease, thus we believe that they likely encapsulate disease status as well.

## Supporting information

Supplementary-Figures_Supplementary-Table-descriptions

## Acknowledgments

This work has been supported by the European Research Council (Synergy Grant DECODE under grant agreement no. 810296) and by the Deutsche Forschungsgemeinschaft (DFG, German Research Foundation) – grant number 433034324. The authors are grateful to M.J. Bonder, L. Marconato, V. Kleshchevnikov, T. Bechtler, H. Öztürk, K. Mikulik, S. Schrod and A. Claringbould for useful discussions and feedback on this work. The authors would also like to thank S. Yazar and J.E. Powell for answering questions regarding the OneK1K dataset. Figure 4h was created with BioRender: Vagiaki, D. (2026) https://BioRender.com/0kpkxr0.

## Author Contributions

*LIVI* was conceptualized by D.V. and M.S. with contributions from T.H. The *LIVI* model and analysis framework was implemented by D.V. and T.H. D.V. performed all the analyses except for the scITD benchmark, which was performed by M.S. O.S. supervised the work. D.V., B.C. and O.S. wrote the manuscript. T.H. and M.S. contributed edits to the manuscript.

## Data availability

All data used for the analysis of this manuscript are publicly available. OneK1K single-cell gene expression and genotype data are available via Gene Expression Omnibus (GSE196830). The gene expression data are also available at https://cellxgene.cziscience.com/collections/dde06e0f-ab3b-46be-96a2-a8082383c4a1. eQTLGen *trans*-eQTL summary statistics are available at https://eqtlgen.org/trans-eqtls.html.

## Code availability

*LIVI* is an open source project available on GitHub: https://github.com/PMBio/LIVI. Code to replicate the analysis in this paper is available on https://github.com/danaivagiaki/LIVI_analyses.

## Methods

### The LIVI framework

#### Background

LIVI is a form of variational autoencoder (VAE) [22, 23], a probabilistic model that uses artificial neural networks to approximate a generative model. Here, we give a brief overview of the main concepts behind VAEs.

Let **x** ∈ ℝ^*M*^ be one data sample generated from latent variables **z** ∈ ℝ^*K*^, according to some generative model *p*(**x**|**z**)*p*(**z**), where *p*(**z**) ∼ 𝒩 (**0, I**) is the prior distribution of the latent variables **z**. VAEs use artificial neural networks, termed *decoders*, to approximate *p*_*θ*_(**x**|**z**). The posterior probability *p*_*θ*_(**z**|**x**) is usually intractable. To make inference feasible, VAEs use another artificial neural network, *q*_*ϕ*_(**z**|**x**), termed *encoder* or *recognition model*, to learn a tractable distribution that approximates the true posterior *p*_*θ*_(**z**|**x**). Typically the recognition model *q*_*ϕ*_(**z**|**x**) is chosen to be Gaussian with diagonal covariance. Any distribution *q*_*ϕ*_ can be used to derive a lower bound on the marginal likelihood *p*_*θ*_(**x**) (*ELBO*):

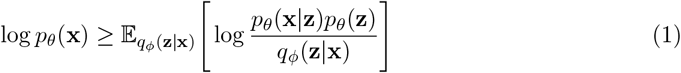

This bound can be optimized with respect to the parameters *ϕ, θ* of the recognition (encoder) and generative (decoder) model, respectively, to maximize the likelihood of the data under the model. The re-parameterization trick [22] enables end-to-end training of the model using stochastic gradient descent.

The ELBO of equation1 can be rewritten as:

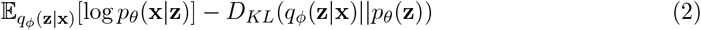

where *D*_*KL*_(*q*_*ϕ*_(**z**|**x**)||*p*_*θ*_(**z**)) is the Kullback-Leibler (KL) divergence between the approximate posterior *q*_*ϕ*_(**z**|**x**) and the prior *p*_*θ*_(**z**) ∼ *N* (**0, I**), a measure of how much these two distributions differ. The term involving *p*_*θ*_(**x**|**z**) is also called the *reconstruction error*, as it represents how well the observed data can be reconstructed given the learned latent representation **z**. Thus, equation 1 becomes:

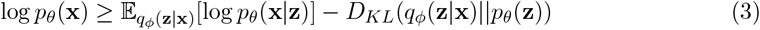

#### The LIVI model

In the population scRNA-seq setting, gene expression profiles for each cell *i* are paired with an donor label *y. LIVI* builds on the basic VAE model and incorporates this grouping structure to capture donor-specific effects on latent gene expression factors. Specifically, *LIVI* takes the vector of observed expression counts for *M* genes, **x**_*i*_ ∈ ℤ^*M*^, of a single-cell *i* from donor *y* as input, and decomposes it into canonical cell state variation, cell state-specific donor effects (*D* × *C*) and global donor effects on gene expression. Cell state variability is encoded into a lower-dimensional *cell-state latent space*, 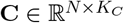, where *N* is the number of cells and *K*_*C*_ is the number of cell-state latent factors. To promote the ability of cell-state latent factors to explain cell state variability, as well as model interpretability, the latent cell-state vector of each cell, **c**_*i*_ ∼ ℝ (**0, I**), is passed through a softmax function to emulate the probability of each cell to belong in a given cell state. Cell state-specific donor effects are learned via a latent linear interaction model between the cell-state factors, **c**_*i*_, and a *cell-state-specific donor factors*, 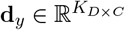, where *K*_*D*×*C*_ is the number of *D* × *C* latent factors:

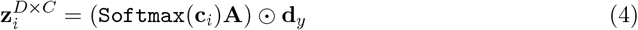

In the above equation, *0* denotes the Hadamard (element-wise) product. The latent variables 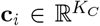 model variation among cells that is shared across all donors, e.g. canonical cell types and states, while the vector **d**_*y*_ represents cell state-specific donor effects on the latent dimensions of **c**_*i*_. 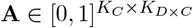 is a factor *assignment matrix* that “assigns” latent factors from 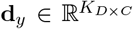 to cell-state latent factors, 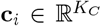 . The values of **A** are enforced to be between 0 and 1 by applying a sigmoid function on a *K*_*C*_ × *K*_*D*×*C*_ matrix of real values, **A**′ (i.e. **A** = *σ*(**A**′)), and pushed towards 0 or 1 by adding a penalty of (**A**(**1** − **A**))^2^ to the ELBO loss (eq.1). This cell-state-specific donor latent space (**Z**_*G*×*C*_) is explicitly decoded via a sparse linear decoder 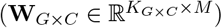 to portray gene regulatory programs. We impose a row-wise *L*_1_ penalty on **W**_*G*×*C*_, so that each *D* × *C* factor has sparse loadings. A second donor embedding, 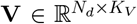, where *N*_*d*_ is the number of donors, is included to account for donor latent variables causing broad changes in gene expression, such as those attributed to population-structure. **V** is decoded by a separate dense linear decoder 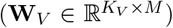. Finally, the reconstructed gene expression vector for cell *i* is given by:

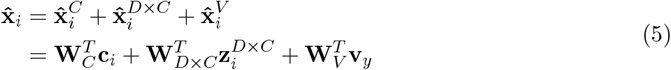

Similar to scVI [16], gene expression profiles are assumed to be drawn from a size-factor-adjusted Negative Binomial distribution, to account for technical noise due to the sampling process and residual over-dispersion:

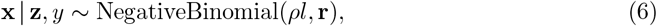

where 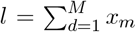 is the total transcript count in a cell, **r** ∈ ℝ^*M*^ is a trainable dispersion parameter for each gene *m*, and *ρ* ∈ [0, 1]^*M*^ represent gene expression frequencies, defined as

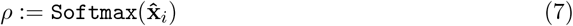

Contrary to scVI, we did not use a zero-inflated negative binomial (ZINB) distribution, as later work showed that the NB distribution can adequately model droplet-based scRNA-seq data [67], since the introduction of unique molecular identifiers (UMIs) in modern droplet-based sequencing technologies eliminates the polymerase chain reaction (PCR) amplification biases that contribute to the observed zero-inflation in early RNA sequencing experiments.

More often than not, scRNA-seq data are generated by different experiments and/or laboratories. These technical confounders are known to influence RNA measurements. When different samples (or donors) are found exclusively in different experimental batches, technical artifacts may mask true genetic effects on gene expression. *LIVI* deals with confounding batch or other covariate effects by learning an *M* -dimensional vector for each confounding covariate (e.g. experimental batch, donor sex, …), **b**_*e*_, which represents the influence of covariate *e* on the expression of each gene *m*. The total effect of *E* covariates on gene expression is the sum of the respective covariate effects:

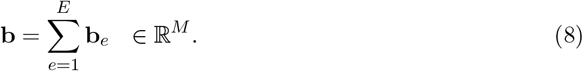

Then equation 5 becomes:

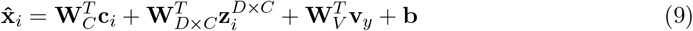

Finally, to increase the power to detect *trans* effects of SNPs, LIVI can explicitly account for known *cis*-eQTLs during training. Assume that among the *M* genes in the dataset, there are *M*_*nocis*_ genes without a known *cis*-eQTL, and *M*_*cis*_ eGenes with a known *cis*-eQTL (where *M*_*nocis*_ + *M*_*cis*_ = *M*), and the lead eSNPs denoted as *G*_*cis*_. LIVI learns an embedding 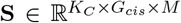, which will add a gene and cell-state-specific *cis* effect correction to the reconstruction term in equation 9, for the lead eSNPs that are present in the current cell (based on the donor genotype). Importantly, the effect of a given eSNP on all other *M* genes besides its corresponding eGene is fixed to 0 across all cell-states.

### Reduced LIVI models

To dissect the utility of the different components of LIVI, we trained three reduced model specifications: 1. a model without the global donor embedding **V** (*LIVI without V*), 2. a model without the *cis*-effect correction (*LIVI without cis*), 3. a model with a single shared decoder for cell-state and donor effects (*LIVI single decoder*), and assessed again the number of fSNPs detected, as well as their replication in eQTLGen [68].

For *LIVI without V*, equation 9 becomes:

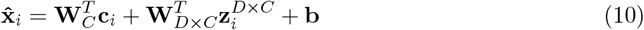

For *LIVI single decoder*, equation 9 becomes:

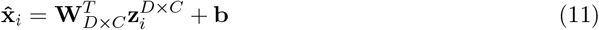

Finally, for *LIVI without cis*, equation 9 remains as is, without the addition of the *cis*-effect correction via 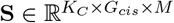.

### Parameter inference

Inference is analogous to the standard VAE. We approximate the true posterior over latent variables **c** using a Gaussian distribution *q*_*ϕ*_(**c**|**x**), where mean and diagonal covariance are parameterized by multi-layer neural networks. The donor embeddings 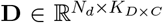 and 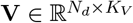, as well as the factor assignment matrix 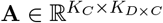, the *cis*-effect embedding 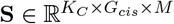 and the covariate correction **b**, are included as part of the observation model and can be optimized using stochastic gradient descent together with the decoder parameters **W**_*C*_, **W**_*D*×*C*_, **W**_*V*_, **r** and the encoder parameters *ϕ*:

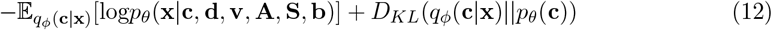

The final model loss includes the sparsity penalties for **W**_*D*×*C*_ (ℒ_*D*×*C*_) and **A** (ℒ_*A*_):

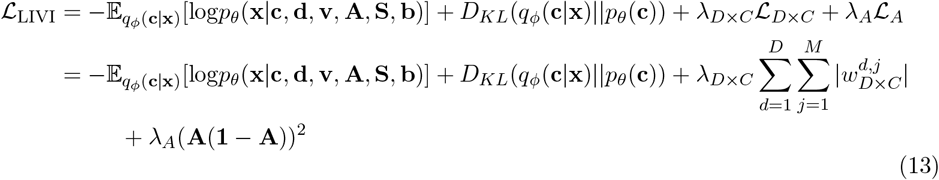

where *λ*_*D*×*C*_ and *λ*_*A*_ control the strength of the penalty for **W**_*D*×*C*_ and **A**, respectively.

LIVI employs a stepwise training scheme, first estimating the cell-state latent factors, **C**, and their loadings, **W**_*C*_. Those parameters are then kept fixed and included in the full model, which is trained end-to-end. By keeping **C** fixed, we eliminate a possible source of non-identifiability, aiding the model to converge to a more stable solution for equation 4.

### Association testing of genetics or other donor characteristics

The matrices 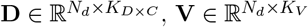 of donor embeddings summarize cell-state-specific and global effects on gene expression factors at the level of donors rather than cells. As a result, they enable fast downstream association testing with donor-level variables of interest. Let 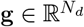 denote a vector of donor variables, e.g., SNP genotypes or clinical phenotypes. To identify interaction effects on factor *k* = 1,…, *K*_*D*×*C*_, one can use a simple linear model for each variable *r*

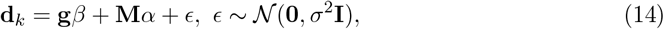

where 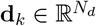, *N*_*d*_ is again the number of donors. In our results, we use a likelihood ratio test for the hypothesis *β* ≠ 0 vs. the null hypothesis *β* = 0. **M** is an optional matrix of confounding effects, e.g., technical batch or genetic population structure. While we have not explored it in this work, an analogous procedure could be used to test for global effects using **V**.

In practice, when 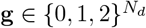 is a vector of SNP genotypes, we use a linear mixed model (LMM) to account for the population structure. For each factor *k*, we model

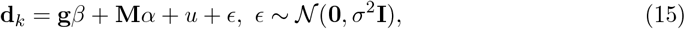

where 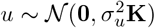 is a random effect parameterized by the kinship matrix between individuals, **K** (computed by PLINK [69]). We used the first 15 principal components (calculated after mean-aggregating gene expression at the individual level) as confounding fixed effects **M**. We fit these LMMs using LIMIX [70]. LIVI’s testing pipeline also includes a multiple testing correction step controlling the false discovery rate (FDR) at a user-specified threshold, which is necessary when testing multiple *D* factors. For our experiments, we used the Benjamini-Hochberg (BH) correction and an FDR of 5% across SNPs and *D* factors. Using the aforementioned procedure, we tested the 9,305 trait-associated SNPs that were previously assessed for *trans* effects as in eQTLGen [25].

Lastly, it should be noted that even if an LMM is used (eq. 15), the testing is done at the donor level (typically *N*_*donors*_ *« N*_*cells*_) and a model is fit for each factor instead for each gene (typically *N*_*factors*_ *« N*_*genes*_), rendering this procedure much more scalable than *CellRegMap* [7]. This testing scheme both alleviates the multiple testing burden of traditional gene-level testing, and better reflects gene regulatory relationships.

### Estimating effect sizes in single cells

While **D** contains donor-level factors, they map to cell-level **D** × **C** factors via equation 4—that is, each **D** factor **d**_*k*_ corresponds to the **D** × **C** factor 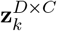. This allows for the association testing to be performed at the level of donors, but the interpretation of the associations at the level of single cells. Specifically, SNP or other donor-level variable effects can be mapped back to the single-cell level by substituting *d*_*y,k*_ with the estimated effect size of SNP *r* on factor **d**_*k*_ (*β*_*r,k*_) in LIVI’s interaction model from equation 4 : *β*_*i,k*_ = (Softmax(**c**_*i*_)**a**_*k*_) *0* **g**_*r*_*β*_*r,k*_, where **a**_*k*_ is the *k*-th column of **A** and **g**_*r*_ is the genotype at SNP *r* in each cell. It is assumed that **g**_*r*_ = **1** to calculate the effect on the cell space, i.e., *in-silico* simulating one allele of the SNP being present in each cell. Then *β*_*i,k*_ is the estimated effect of SNP *r* on cell *i*.

### Assigning *D* factors to known cell types

*D* factors are assigned to cell-state factors that have assignment values ≥ 0.9 in the **A** matrix. Cell-state factors can be mapped to known cell types by averaging the cell-state factor values across cells of each cell type, and subsequently selecting the cell type with the highest mean value (across cells) for the given cell-state factor. However, for visualization purposes, it is often preferable to have a 1:1 correspondence between *D* factors and cell types. In this case, a cell-types × *D* matrix can be calculated, and each *D* factor can be assigned to the cell type (row) that has the maximum mean value (across cells) for that *D* factor (column). This approach utilizes the inherent matrix multiplication taking place during model training (**CA** in equation 4), however, it also forces the discretization of cell states to cell types (and consequently cell type-specific effects), even though the given factor could represent a cellular program characterizing multiple closely related cell types.

### Determining the number of genes explaining *D* × *C* factors

*D* × *C* factors are sparse, with only a small subset of the genes having clearly distinguishable large contributions to them. To ensure only those genes are selected when employing an automated, data-driven selection strict thresholds should be used. To this end, for each *D* × *C* factor we selected genes with loadings more than 100 times above or below the interquartile range (IQR).

### Correlation of SNP effects at the single-cell level across different stochastic model runs

To investigate the concordance of the predicted SNP effect at the single-cell-level across different model initializations, we ran *LIVI* with five different random seeds and the same set of hyperparameters. Subsequently, we performed association testing with the same set of SNPs. First, we counted the number of runs that discovered each fSNP. Then we selected a subset of fSNPs that were discovered across all five runs and were significantly associated with only one *D* factor in each run. For those fSNPs, we calculated their effect on the cell space (see section *Association testing of genetics or other donor characteristics*) for each run, and then we computed pairwise Pearson correlations of the estimated cell-level effect between any two runs, as well as the mean across all pairwise Pearson’s correlations. Restricting to fSNPs that were associated with one *D* factor only was a necessary simplification in order to make the above analyses computationally tractable. Moreover, the variance explained by a given *D* × *C* factor in one run might be explained by two or more *D* × *C* factors in another run. However, *D* factors uniquely associated with SNPs discovered consistently across runs are more likely to explain the same variation in the data, and thus simpler to compare.

### Data processing

#### OneK1K

##### Genotype data

We filtered the accurately imputed (INFO *>* 0.8, yielding 12,108,283 variants; see original publication for details) genotypes of [8] for minor allele frequencies (MAF) ≥ 0.01 and Hardy-Weinberg equilibrium (HWE) *p*-value *>* 10^−6^ using PLINK 1.9 [69], giving a final number of 7,405,854 variants. Finally, from those variants, we selected 9,305 trait-associated SNPs that were also assessed for *trans* effects in eQTLGen [25].

##### scRNA-seq

We used the raw count scRNA-seq data from [8]. Starting with 1,272,489 cells and 32,738 genes, we removed cells in which fewer than 500 genes were expressed (95,576 cells removed), and genes not expressed in any cells (4,580 genes removed). Raw count data were then normalized according to the median of total counts across cells, and log-transformed (ln(*x* + 1)) using scanpy [71]. We further excluded non-immune cells, namely platelets and erythrocytes, from any analysis. Normalized counts were used to select 10,000 highly variable genes (HVGs) with scanpy (`sc.pp.highly variable genes(adata, n top genes=10000, flavor=“seurat”`)) and the 10,000 highest expressed genes. We used the union of those genes to train *LIVI*. After these filters, *LIVI* was trained on raw count data of 14,212 genes, measured in 1,172,790 immune cells.

##### eQTLGen

To assess the power of *LIVI* to discover genetic associations, we tested the same trait-associated SNPs as in eQTLGen, a large bulk meta-analysis study of 31,684 samples [25], for effects on *LIVI* ‘s donor factors. 10,317 trait-associated SNPs were assessed for *trans*-eQTL effects in eQTLGen (*trans*-eQTLGen). We tested the 9,305 of those that were present in the OneK1K cohort. Out of the 9,305 variants, 2,581 showed significant *trans* effects in eQTLGen. We compared the number of discoveries obtained by *LIVI* to that of the original study, as well as to those obtained by alternative methods (see section *Benchmarks*) on the same 9,305 variants and 14,212 genes. In order to assess the extent to which LIVI captures *cis* vs. *trans* effects, we further compared our discoveries with *cis*-eQTLGen [25]. For a fair comparison, we only took into account the 6,405 variants that displayed a significant *cis*-eQTL effect on the 14,212 genes that were measured in the OneK1K dataset and used to train *LIVI*.

### Data analysis

#### Fitting LIVI on the OneK1K data

We trained LIVI on the OneK1K data using 15 cell-state, 700 *D* and 5 *V* factors. We set the weights of the sparsity penalties (*λ*_*D*×*C*_ and *λ*_*A*_ in equation 13) to 10^−3^. We trained the cell-state latent factors **C** and corresponding **W**_*C*_ decoder for 60 epochs and subsequently froze them. The rest of the model components were trained until convergence (317 epochs in total) with a batch size of 1024. In general, we recommend pre-training of the cell-state factors and decoder for 20-60 epochs and training the donor-related components of the model for a minimum of 60 epochs.

#### Factor annotation using gene set enrichment analysis

All gene set enrichment analyses (GSEA) were performed using Enrichr [72] as implemented in the gseapy [73] package (version 1.1.4), with a different set of selected genes and databases as described in the subsections below. For all analyses, we used all 14,212 genes that *LIVI* was trained on as the background set.

#### Annotation of cell-state factors

For each cell-state factor, we selected genes with loadings greater than 0.5× the interquartile range (IQR) for that factor. We used those marker genes to query the Azimuth (as of 2021) cell type annotation database and obtain known cell type labels. First we selected all the terms (cell type annotations) with adjusted *p*-value ≤ 0.05, and finally the five terms with the smallest adjusted *p*-values to annotate a given factor. Cells with high loadings for the specific factor got assigned to the corresponding cell type(s).

#### Annotation of *D* × *C* factors

We performed GSEA using the top genes for each *D* × *C* factor associated with at least one SNP. Top genes were selected as described in section *Determining the number of genes explaining D* × *C factors*. The analysis was run separately for four databases: Kyoto Encyclopedia of Genes and Genomes (KEGG) (as of 2021), Reactome (as of 2022), Gene Ontology (GO) Biological Process (as of 2023) and GO Molecular Function (as of 2023). For each database, we retained terms with an adjusted *p*-value ≤ 0.05. When multiple terms were significantly enriched within a database, we selected the term involving the largest number of genes; if several terms involved an equal number of genes, we chose the one with the smallest adjusted *p*-value and highest odds ratio. For visualization purposes, we considered only highly variable *D* × *C* factors, defined as those with variance across cell types greater than the mean factor variance across cell types.

#### Polygenic (risk) score calculation

We tested seven autoimmune diseases and twelve blood traits for associations with LIVI factors: inflammatory bowel disease (IBD) and ulcerative colitis (UC), rheumatoid arthritis (RA), type 1 diabetes (T1D), multiple sclerosis (MS), celiac disease (CeD), psoriasis, as well as monocyte count and percentage, lymphocyte count and percentage, leukocyte count, red blood cell count (RBC) and distribution width (RBDW), platelet count and distribution width (PDW), plateletcrit, mean platelet volume (MPV), mean corpuscular volume (MCV). PRSs for each disease/trait were calculated using the PGS Catalog [43] pipeline (https://github.com/pgscatalog/pgsc_calc). We used precomputed score summary statistics available in the PGS Catalog as follows: PGS000017 for IBD, PGS002066 for UC, PGS002088 for RA, PGS002025 for T1D, PGS002038 for MS, PGS002067 for CeD, PGS002083 for Psoriasis, PGS002186 for monocyte count, PGS002208 for monocyte percentage, PGS002183 for lymphocyte count, PGS002203 for lymphocyte percentage, PGS002180 for leukocyte count, PGS002123 for RBC, PGS002122 for RBDW, PGS002188 for plateletcrit, PGS002191 for platelet count, PGS002190 for PDW, PGS002189 for MPV, and PGS002207 for MCV. We also tested for associations with PGS for height (PGS002146), myopia (PGS002211) and hair color (PGS002109) as negative controls, i.e., we were expecting no associations between height, myopia or hair color and LIVI factors. Prior to association testing, PRS/PGS values for a given disease/trait were standardized to zero mean and unit variance.

### MEME/FIMO analysis

ZFP57 binding consensus sequence was downloaded from JASPAR (https://jaspar.elixir.no/; 2024 version) [75], and *KLRB1* sequence was downloaded from UCSC Genome Browser (https://genome.ucsc.edu/index.html). *KLRB1* sequence was scanned for ZFP57 motif occurrences using FIMO (https://meme-suite.org/meme/tools/fimo) [32]) with the default p-value threshold of 1*e*^−4^.

#### Assessment of continuous effects

To assess the extent to which LIVI captures genetic effects acting beyond discrete cell types, we considered fSNPs associated with *D* factors that map to cell-state factors with high activity in the following cellular differentiation trajectories: myeloid lineage (cell-state factor 6), naive B cell to plasma cell (cell-state factor 7), naive/central memory to effector memory CD4+ T cells (cell-state factor 5), and naive/central memory to effector memory CD8+ T cells (cell-state factor 11). Following this approach, we found that:

- 12 *D* factors mapped to cell-state factor 6. These *D* factors were associated with 56 SNPs, 36 out of which were not detectable by single-gene testing.
- 22 *D* factors mapped to cell-state factor 7. These *D* factors were associated with 288 SNPs, 189 out of which were not detectable by single-gene testing.
- 10 *D* factors mapped to cell-state factor 5. These *D* factors were associated with 188 SNPs, 126 out of which were not detectable by single-gene testing.
- 13 *D* factors mapped to cell-state factor 11. These *D* factors were associated with 76 SNPs, 51 out of which were not detectable by single-gene testing.

### Benchmarks

#### Principal component analysis

Principal components (PCs) capture gene expression variability, and even though the variance captured by the first PCs is generally attributable to global variables, such as technical batch, an increasing number of PCs is likely to capture smaller sources of variation, such as SNP effects. To mimic *LIVI* ‘s individual embedding, we calculated the mean gene expression for each donor across all cells belonging to the same pre-annotated cell type from [8] (“CD4 ET”, “CD4 SOX4”, “CD4 NC”, “CD8 S100B”, “CD8 ET”, “B IN”, “CD8 NC”, “B Mem”, “NK”, “NK R”, “Mono NC”, “Mono C”). PCA was performed on these pseudobulk donor-level samples (i.e. within each cell type). This approach gives a *cell types* × *donors* × *PCs* embedding, which can be easily tested for genetic associations using *LIVI* ‘s testing pipeline (see section *Association testing of genetics or other donor characteristics*). We tested the same number of PCs as the number of *D* factors, *K*_*D*×*C*_.

#### scITD

To benchmark scITD [20] against LIVI, we applied the scITD method to the OneK1K scRNA-seq data and tested factor–genotype associations using LIVI’s testing pipeline (see *Association testing of genetics or other donor characteristics*). scITD constructs donor-by-cell-type pseudobulk expression tensors and performs Tucker tensor decomposition to learn shared axes of variation across cell types. We generated two separate tensors: one at a coarse-grained (L1) cell type level using five broad PBMC types (“B”, “CD4T”, “CD8T”, “Mono”, “NK”), including donors with at least two cells per cell type (n = 960); and one at a fine-grained (L2) level using twelve cell types as annotated in [8] (“CD4 ET”, “CD4 SOX4”, “CD4 NC”, “CD8 S100B”, “CD8 ET”, “B IN”, “CD8 NC”, “B Mem”, “NK”, “NK R”, “Mono NC”, “Mono C”), including donors with at least one cell per cell type (n = 549). Since the fine-grained L2-level approach came at the expense of losing approximately 56% of the samples (549 donors instead of 981), we proceeded with the L1-aggregated tensor. We trained scITD five times using different numbers of factors (100, 300, 500, 700 and 900 factors). Subsequently, we tested the donor loadings for each factor for association with the same 9,305 trait-associated variants that were also used for testing LIVI’s factors (see section *eQTLGen*). This allowed for the direct comparison of the number fQTLs detected by scITD-derived factors vs. LIVI-derived factors.

#### MrVI

We compared *LIVI* ‘s ability to discover genetic associations to that of MrVI [21], another VAE-based model that uses raw count scRNA-seq data from multiple individuals to capture the effect of donor characteristics on gene expression. To this end, we trained MrVI with the same number of cell-state (corresponding to *K*_*C*_) and donor-informed (corresponding to *K*_*D*×*C*_) latent dimensions as LIVI. Next, we aggregated the resulting donor-aware cell latent space (referred to as *z* in [21]) at the donor level, by taking the mean of values for cells belonging to the same individual. We used the resulting **z**_**aggr**_ embedding as input to LIVI’s association testing framework to assess the effects of the same 9,305 trait-associated variants.

#### Single-gene testing

Traditional gene-level *trans*-eQTL mapping within each cell type was performed using tensorQTL version 1.0.10 [76, 77]. We considered the same set of genes used to train LIVI (see section *OneK1K*). For each gene, we tested the subset of the 9,305 genetic variants from *trans*-eQTLGen (see section *eQTLGen*) that were located either on a different chromosome or further than 5Mb away on the same chromosome. Finally, multiple testing correction was performed in different ways, depending on the analysis task. When comparing the power of latent variable methods against the power of gene-level testing to discover eQTLs, multiple testing correction was done in two steps to account for the vast difference in the number of tests performed by the respective approaches. To this end, we first performed Bonferroni correction for the number of phenotypes tested, i.e., the number of latent factors for LIVI and scITD (*n* = 700), the number of PCs × number of cell types for PCA (*n* = 700 ∗ 14 = 9800), and finally the number of genes × number of cell types (*n* = 14, 212 ∗ 14 = 198, 968) for gene-level testing. Second, we performed Benjamini-Hochberg (BH) correction to control for FDR < 5%. For a qualitative comparison between LIVI and gene-level testing, i.e. *which* SNPs were identified by each approach, we followed the scheme typically applied in eQTL studies and controlled only for FDR < 5%.

