## Supplementary-Figures_Supplementary-Table-descriptions for "Mapping *trans*-eQTLs at single-cell resolution using Latent Interaction Variational Inference"

D. Vagiaki *et al.*

### Supplementary Tables

Supplementary Tables S2 to S6 can be found online in Microsoft Excel format, file name "Supplementary_Tables_S2-S6.xlsx". Supplementary Tables S7 to S11 can be found online in Microsoft Excel format, file name "Supplementary_Tables_S7-S11.xlsx". All Supplementary Table descriptions are provided below.

**Table S1. Conceptual comparison of LIVI to other latent variable methods. Method:** Method name, **Reference**: Reference to the peer-reviewed publication introducing the method, **Single-cell expression counts**: Accepts single-cell expression counts as input, **Preannotated cell types**: Requires preannotated cell types, **Generic donor representation**: Learns a generic donor representation, uninformed of specific donor covariates, **Genotypes during training**: Uses genotypes during parameter estimation, **Separate cell-state and donor/genetic latent representation**: Learns separate latent representations for cell-states vs. donors/genetics, **Separate cell-state and donor/genetic gene programs**: Learns separate gene programs to explain cell-state vs. donor/genetic effects, **Designed for genetics:** Designed to detect genetic effects on gene expression, **Focus on *cis* vs. *trans* effects:** If designed for genetics, if the focus is on *cis* or *trans* effects, **Single-gene tests:** Employs single-gene association tests to identify genetic variant/donor effects on gene expression.

| **Method** | **LIVI** | **MrVI** | **scITD** | **SURGE** |
|---|---|---|---|---|
| **Reference** | — | Boyeau 2025, *Nat Methods* | Mitchel 2024, *Nat Biotechnol* | Strober 2024, *Genome Biol* |
| **Single-cell expression counts** | Yes | Yes | No | No |
| **Preannotated cell types** | No | No | Yes | Yes |
| **Generic donor representation** | Yes | Yes | Yes | No |
| **Genotypes during training** | No | No | No | Yes |
| **Separate cell-state and donor/genetic latent representation** | Yes | Yes | No | No |
| **Separate cell-state and donor/genetic gene programs** | Yes | No | No | No |
| **Designed for genetics** | Yes | No | No | Yes |
| **Focus on *cis* vs. *trans* effects** | *trans* | — | — | *cis* |
| **Single-gene tests** | No | No | No | Yes |

**Table S2. Top genes for each $D \times C$ factor, defined as those with absolute loadings greater than $100 \times$IQR.**

| Column | Description |
|---|---|
| Factor | Factor ID |
| TopGenes | Semicolon-separated list of HGNC gene symbols |

**Table S3. Annotation of $D \times C$ factors using GSEA on their top genes.**

| Column | Description |
|---|---|
| Factor | Factor ID |
| Pathway | Enriched term |
| Genes | Factor genes involved in the enriched term |
| P-value | $p$-value |
| Adjusted P-value | Adjusted $p$-value |
| Odds Ratio | Measure of the over-representation of genes in the particular enriched term |
| Gene_set | Database source: KEGG or Reactome or Gene Ontology (GO) |

**Table S4. Output summary statistics of discovered fQTLs using LIVI (FDR $< 5\%$).**

| Column | Description |
|---|---|
| Factor | Factor ID |
| SNP_id | SNP ID based on hg19 genomic coordinates |
| effect_size | Effect size coefficient |
| effect_size_se | Effect size coefficient standard error |
| p_value | $p$-value |
| assessed_allele | SNP allele used |
| corrected_pvalue | $p$-value adjusted using Benjamini-Hochberg correction |

**Table S5. Output summary statistics of significant single-gene SNP associations (FDR $< 5\%$).**

| **Column** | **Description** |
|---|---|
| SNP_id | SNP ID based on hg19 genomic coordinates |
| gene | ENSEMBL gene ID |
| p_value | $p$-value |
| effect_size | Effect size coefficient |
| effect_size_se | Effect size coefficient standard error |
| ref_allele | Reference allele |
| alt_allele | Alternative allele assessed for effect on gene expression |
| celltype | Cell type in which the expression of the gene was measured |
| Storey_q | $p$-value adjusted using Storey-Q method |
| BH_corrected_pvalue | $p$-value adjusted using Benjamini-Hochberg |
| BY_corrected_pvalue | $p$-value adjusted using Benjamini-Yekutieli |

**Table S6. $P$-values of all single-gene associations with rs1610677.**

| **Column** | **Description** |
|---|---|
| SNP_id | SNP ID based on hg19 genomic coordinates |
| gene | ENSEMBL gene ID |
| p_value | $p$-value |
| ref_allele | Reference allele |
| alt_allele | Alternative allele assessed for effect on gene expression |
| celltype | Cell type in which the expression of the gene was measured |
| eQTL | eQTL ID defines as <SNP_id>__<gene>__<celltype> |

**Table S7. $P$-values of all single-gene associations with rs61907765.**

| Column | Description |
|---|---|
| SNP_id | SNP ID based on hg19 genomic coordinates |
| gene | ENSEMBL gene ID |
| p_value | $p$-value |
| ref_allele | Reference allele |
| alt_allele | Alternative allele assessed for effect on gene expression |
| celltype | Cell type in which the expression of the gene was measured |
| eQTL | eQTL ID defines as <SNP_id>__<gene>__<celltype> |

**Table S8. $P$-values of all single-gene associations with rs12550612.**

| Column | Description |
|---|---|
| SNP_id | SNP ID based on hg19 genomic coordinates |
| gene | ENSEMBL gene ID |
| p_value | $p$-value |
| ref_allele | Reference allele |
| alt_allele | Alternative allele assessed for effect on gene expression |
| celltype | Cell type in which the expression of the gene was measured |
| eQTL | eQTL ID defines as <SNP_id>__<gene>__<celltype> |

**Table S9. Output summary statistics of significant PRS-LIVI factor associations (FDR $< 5\%$).**

| Column | Description |
|---|---|
| Factor | Factor ID |
| PRS | Polygenic (risk) score |
| effect_size | Effect size coefficient |
| effect_size_se | Effect size coefficient standard error |
| p_value | $p$-value |
| assessed_allele | SNP allele used |
| corrected_pvalue | $p$-value adjusted using Benjamini-Hochberg correction |

**Table S10. Mapping of associations between LIVI factors and PRSs for autoimmune diseases.**

| Column | Description |
|---|---|
| PRS_group | Group of disease(s) associated with the same $D$ factors |
| Factor | $D$ Factor ID |
| PRS | Polygenic Risk Score |
| effect_size | Effect size coefficient |
| effect_size_se | Effect size coefficient standard error |
| p_value | $p$-value |
| corrected_pvalue | $p$-value adjusted using Benjamini-Hochberg correction |
| TopGenes | Semicolon-separated list of HGNC symbols of genes estimated to explain the associated $D$ factor |
| Celltype | Known cell type assigned to the associated $D$ factor |

**Table S11. Output summary statistics of significant single-gene PRS association (FDR $< 5\%$).**

| Column | Description |
|---|---|
| PRS | Polygenic (risk) score |
| gene | ENSEMBL gene ID |
| p_value | $p$-value |
| effect_size | Effect size coefficient |
| effect_size_se | Effect size coefficient standard error |
| celltype | Cell type in which the expression of the gene was measured |
| Storey_q | $p$-value adjusted using Storey-Q method |
| BH_corrected_pvalue | $p$-value adjusted using Benjamini-Hochberg |
| BY_corrected_pvalue | $p$-value adjusted using Benjamini-Yekutieli |

### Supplementary Figures

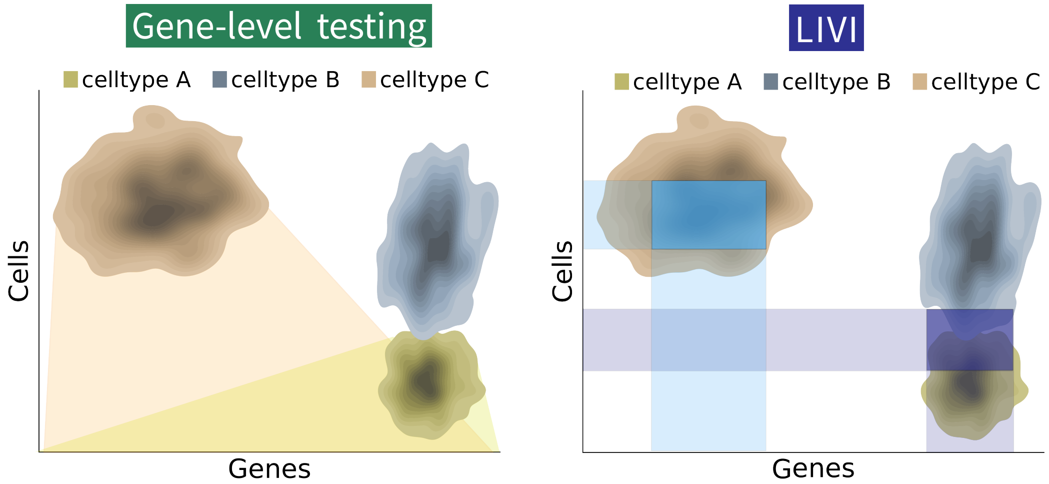

**Figure S1. Illustration of conceptual differences between conventional eQTL mapping *vs.* eQTL mapping using a single-cell latent variable model such as LIVI.** Conventional eQTL mapping strategies consider the expression profile of individual genes, aggregated at the level of discrete cell types (left). Pseudobulk profiles of individual genes in each discrete cell type are then used as input for eQTL mapping. By contrast, LIVI estimates $D \times C$ factors, which capture interindividual variation of sets of genes in subsets of cells, defined in a data-driven manner (right). Thus, $D \times C$ factors may capture variation in cell populations that span related cells types or cell type subsets, defining a distinct cellular program.

**a**

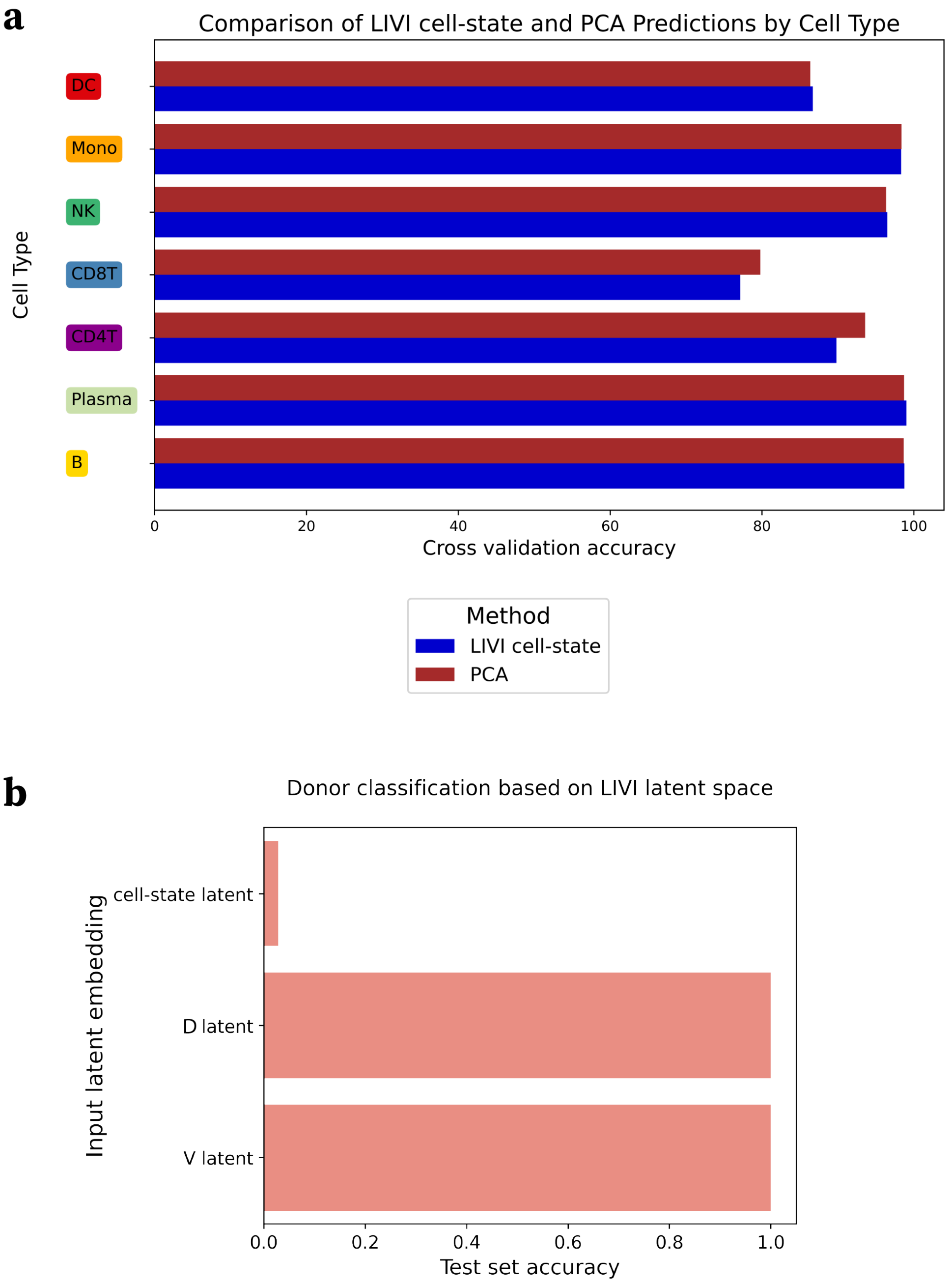

**b**

**Figure S2. Assessment of the disentanglement of cell-state and donor variation in LIVI.** **(a)** Cell type classification accuracy based on logistic regression estimators, using LIVI's cell-state latent factors or an equal number of PCs (15) calculated on the same dataset as input. The experiment was repeated five times, each time using a different subset of the dataset for training and testing (5-fold-cross-validation). The average cross-validation accuracy is reported. **(b)** Donor classification accuracy using the different LIVI latent embeddings ($\mathbf{C}, \mathbf{D}, \mathbf{V}$) as input to a feed-forward neural network (FNN). 90% of the dataset was used to train the FNN, while the remaining 10% of unseen data (test set) was used to assess the accuracy.

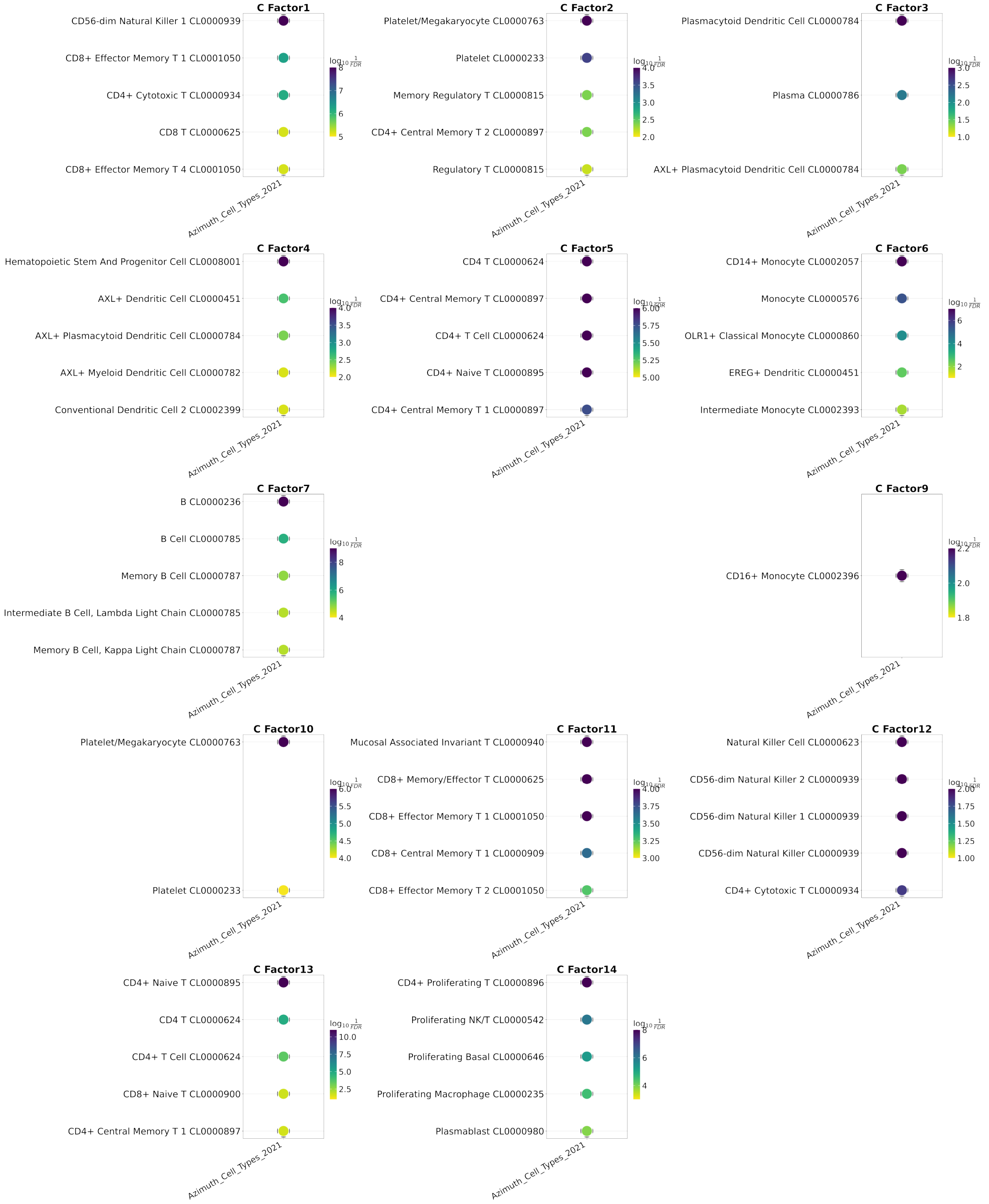

**Figure S3. LIVI cell state factors capture canonical cell type marker genes.** In case of datasets without *a priori* known cell types, LIVI's cell-state factors can be used to annotate cells based on known cell types from publicly available databases. Specifically, genes with high loadings for each cell-state factor can be utilized in a gene set enrichment analysis (GSEA) framework to query databases, such as Azimuth, for cell type marker genes. After mapping cell-state factors to known cell types, cells with high values for a given factor are assigned to its corresponding cell type. Here we illustrate the GSEA terms for the 15 cell-state factors used to train LIVI on the OneK1K dataset. Factors 8 and 15 are omitted, as they did not yield any significant enrichment against the Azimuth database.

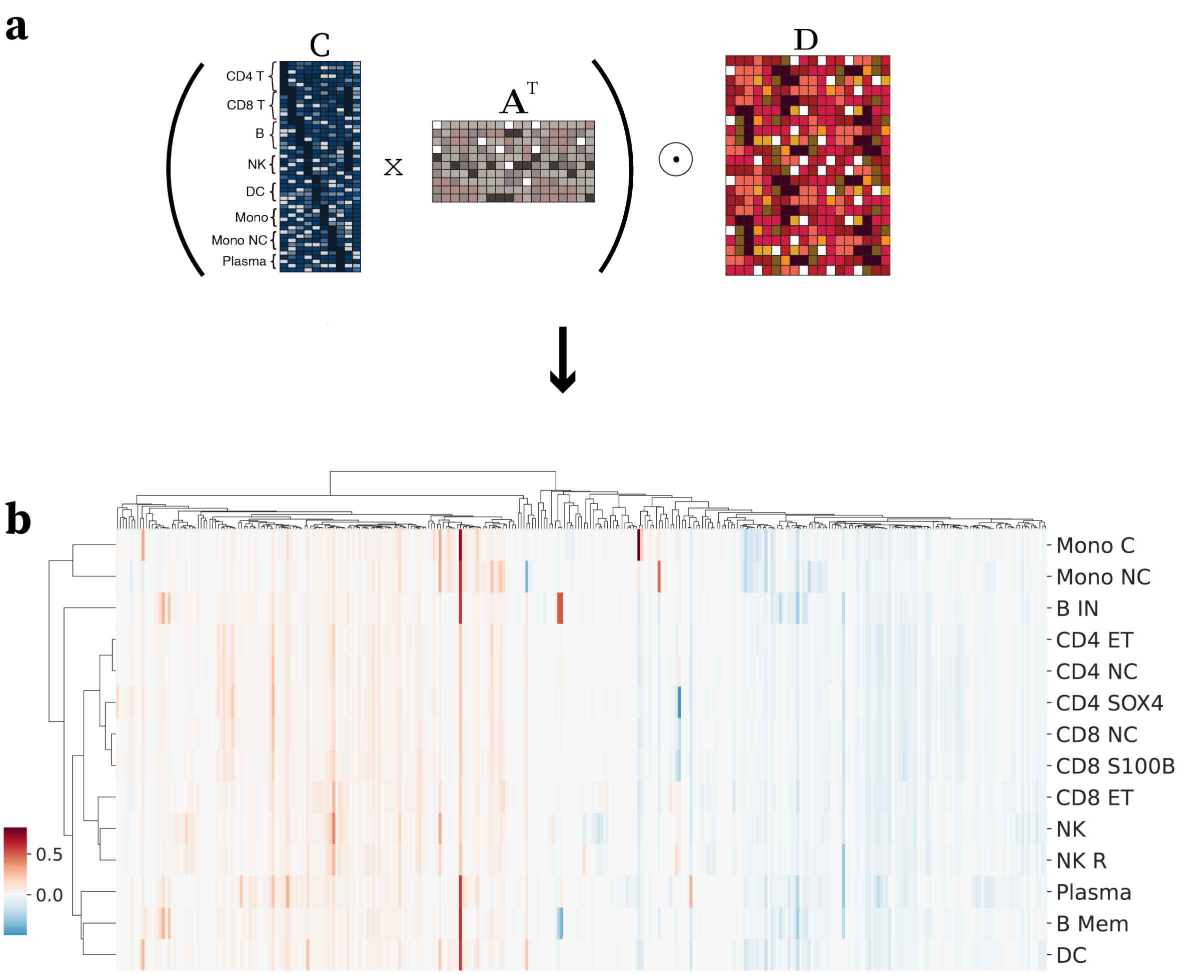

**Figure S4. Identification of LIVI's $D \times C$ factors and activity across cell types. (a)** Illustration of LIVI's latent interaction model for a scRNA-seq dataset of $N$ cells: $\mathbf{D} \in \mathbb{R}^{N \times K_{D \times C}}$ factors (red matrix), which capture interindividual variation, are mapped to cell-state factors, $\mathbf{C} \in \mathbb{R}^{N \times K_C}$ (blue matrix), which capture cell state variation, via the assignment matrix $\mathbf{A} \in [0, 1]^{K_C \times K_{D \times C}}$ (grey matrix). The element-wise ($\odot$) product between $\mathbf{CA}$ and $\mathbf{D}$ encompasses cell-state-specific donor effects ($\mathbf{D} \times \mathbf{C} \in \mathbb{R}^{N \times K_{D \times C}}$). Note that the $\mathbf{D}$ donor factors have been expanded to the cell level by using for each cell the row $\mathbf{d}$ that corresponds to the latent representation of the donor the specific cell originates from. **(b)** $D \times C$ factors associated with at least one genetic variant were averaged across cells of the same annotated cell type from the original publication (Yazar 2022), and subsequently clustered based on cosine similarity.

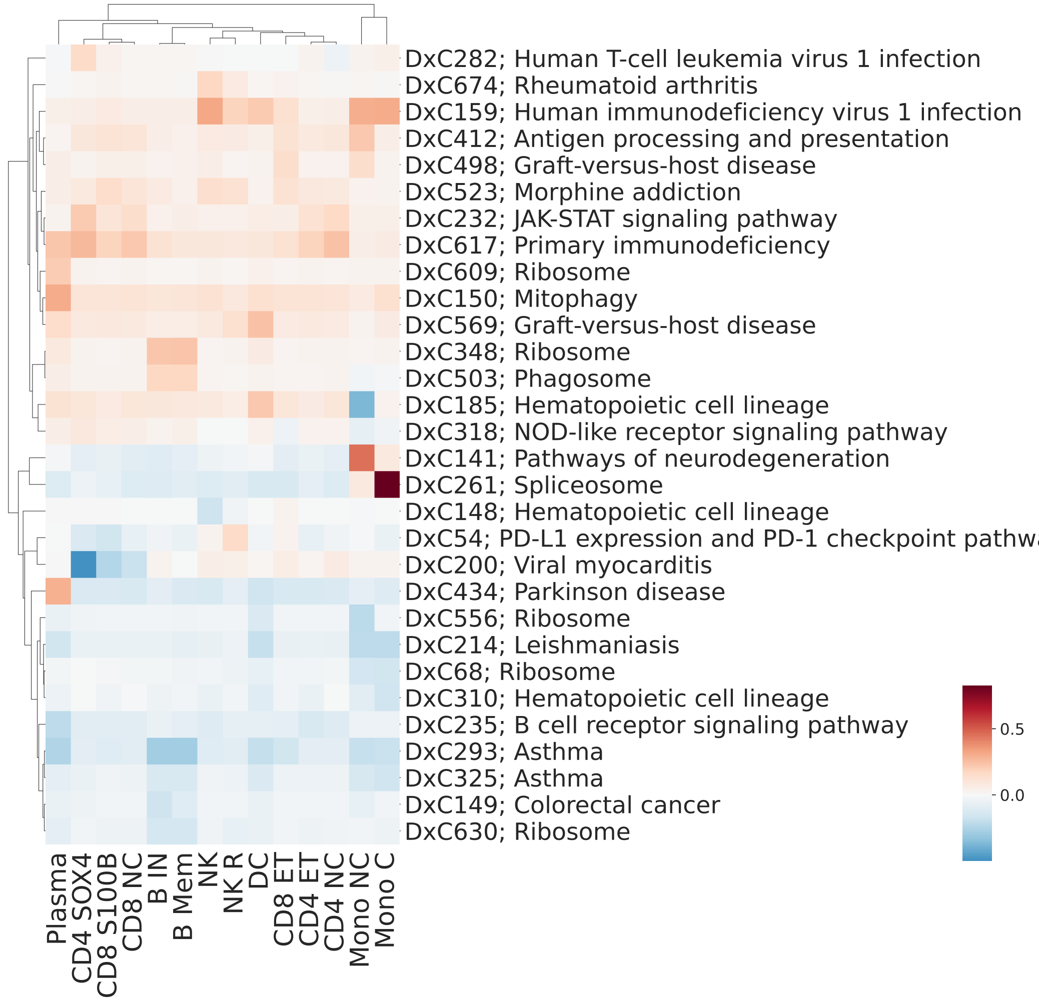

**Figure S5. Annotations of $D \times C$ factors based on KEGG database.** For visual clarity, we considered only highly variable $D \times C$ factors, defined as those with variance across cell types greater than the mean factor variance across cell types. For each of those highly variable $D \times C$ factors, genes with loadings greater or smaller than $100 \times$ IQR were assessed for enrichment of specific biological pathways. Among the significantly enriched terms for each factor, we selected the one involving the largest number of genes; if this did not yield a unique term, we chose the one with the smallest adjusted $p$-value; if this still did not yield a unique term, we chose the one with the highest odds ratio (among the terms with equal numbers of genes and adjusted $p$-values). For visualization purposes, we considered only highly variable $D \times C$ factors, defined as those with variance across cell types greater than the mean factor variance across cell types.

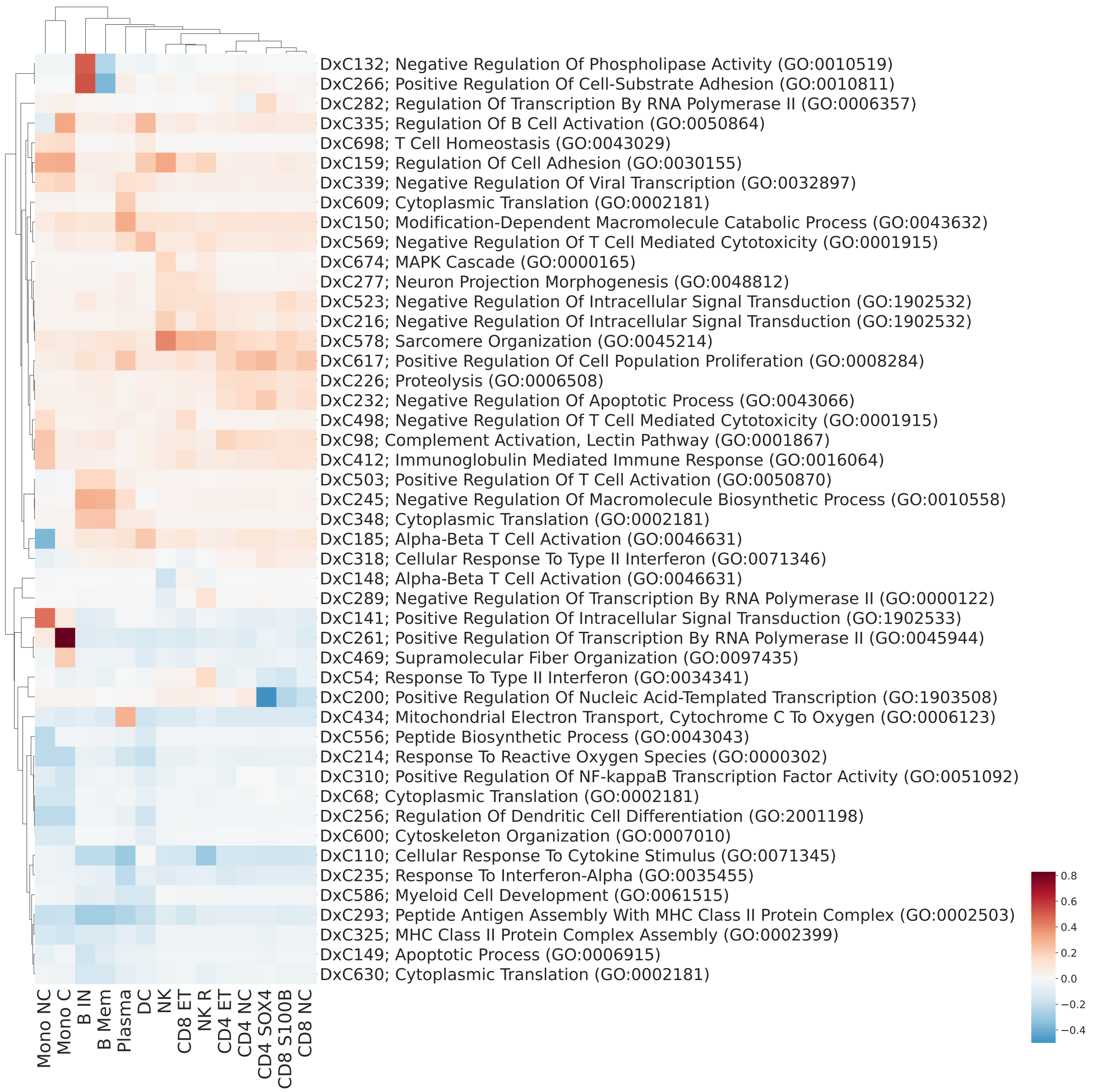

**Figure S6. Annotations of $D \times C$ factors based on Gene Ontology Biological Process database.** For visual clarity, we considered only highly variable $D \times C$ factors, defined as those with variance across cell types greater than the mean factor variance across cell types. For each of those highly variable $D \times C$ factors, genes with loadings greater or smaller than $100 \times$ IQR were assessed for enrichment of specific biological pathways. Among the significantly enriched terms for each factor, we selected the one involving the largest number of genes; if this did not yield a unique term, we chose the one with the smallest adjusted $p$-value; if this still did not yield a unique term, we chose the one with the highest odds ratio (among the terms with equal numbers of genes and adjusted $p$-values).

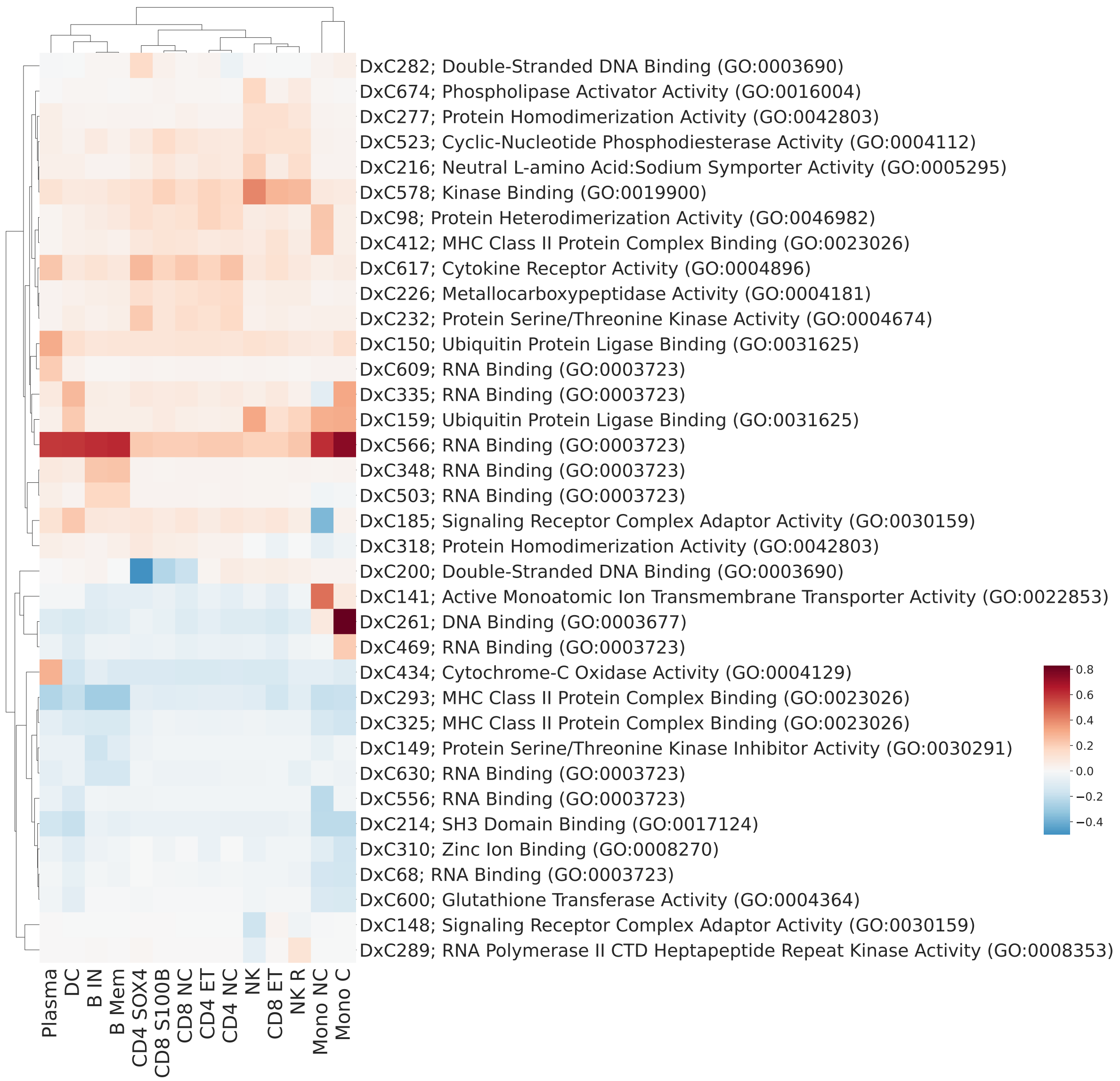

**Figure S7. Annotations of $D \times C$ factors based on Gene Ontology Molecular Function database.** For visual clarity, we considered only highly variable $D \times C$ factors, defined as those with variance across cell types greater than the mean factor variance across cell types. For each of those highly variable $D \times C$ factors, genes with loadings greater or smaller than $100 \times$ IQR were assessed for enrichment of specific biological pathways. Among the significantly enriched terms for each factor, we selected the one involving the largest number of genes; if this did not yield a unique term, we chose the one with the smallest adjusted $p$-value; if this still did not yield a unique term, we chose the one with the highest odds ratio (among the terms with equal numbers of genes and adjusted $p$-values).

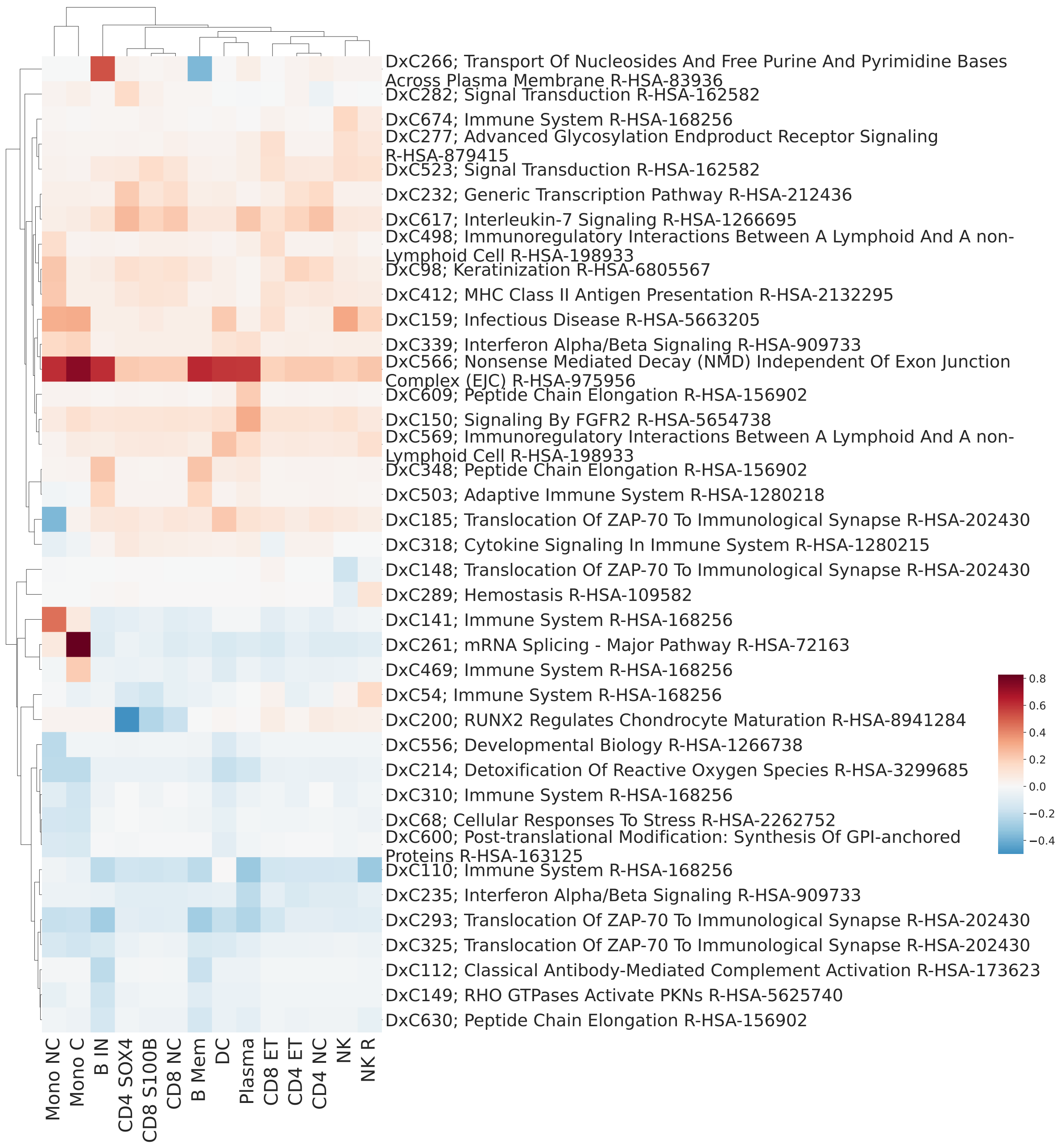

**Figure S8. Annotations of $D \times C$ factors based on Reactome database.** For visual clarity, we considered only highly variable $D \times C$ factors, defined as those with variance across cell types greater than the mean factor variance across cell types. For each of those highly variable $D \times C$ factors, genes with loadings greater or smaller than $100 \times$ IQR were assessed for enrichment of specific biological pathways. Among the significantly enriched terms for each factor, we selected the one involving the largest number of genes; if this did not yield a unique term, we chose the one with the smallest adjusted $p$-value; if this still did not yield a unique term, we chose the one with the highest odds ratio (among the terms with equal numbers of genes and adjusted $p$-values).

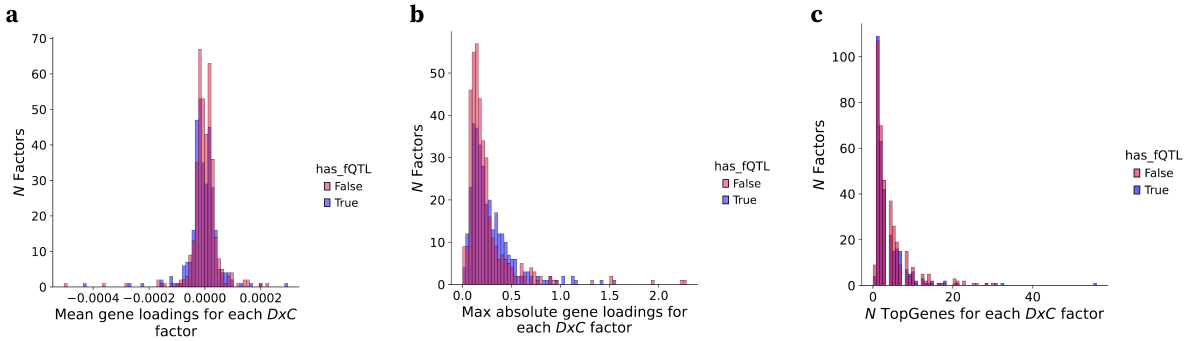

**Figure S9. Sparsity of $D \times C$ factors.** **(a)** Histogram of mean loadings for $D \times C$ factor. Color denotes whether a given $D \times C$ factor was significantly associated with at least one SNP (`has_fQTL = True`; FDR $< 5\%$). **(b)** Histogram of maximum absolute loadings for $D \times C$ factors. Color code as in **(a)**. **(c)** Histogram of number of genes explaining each $D \times C$ factor (Methods). Color code as in **(a)**.

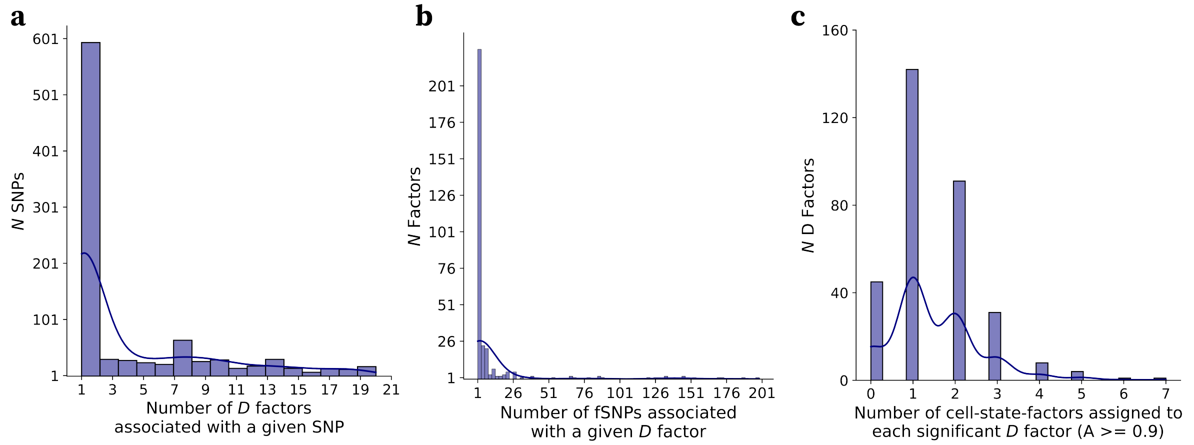

**Figure S10. LIVI identifies cell state-specific genetic effects.** **(a)** Number of unique $D$ factors a given *fSNP* is associated with. **(b)** Number of unique *fSNPs* a given $D$ factor is associated with. **(c)** Number of cell-state factors assigned to each *fSNP*-associated $D$ factor $k$ (i.e. rows of the $\mathbf{A}$ matrix with values $\geq 0.9$, $\mathbf{a}_{\mathbf{d_k}} \geq 0.9$). Most *fSNP*-associated $D$ factors are assigned to a single cell-state factor, implying that the associations are cell-state-specific.

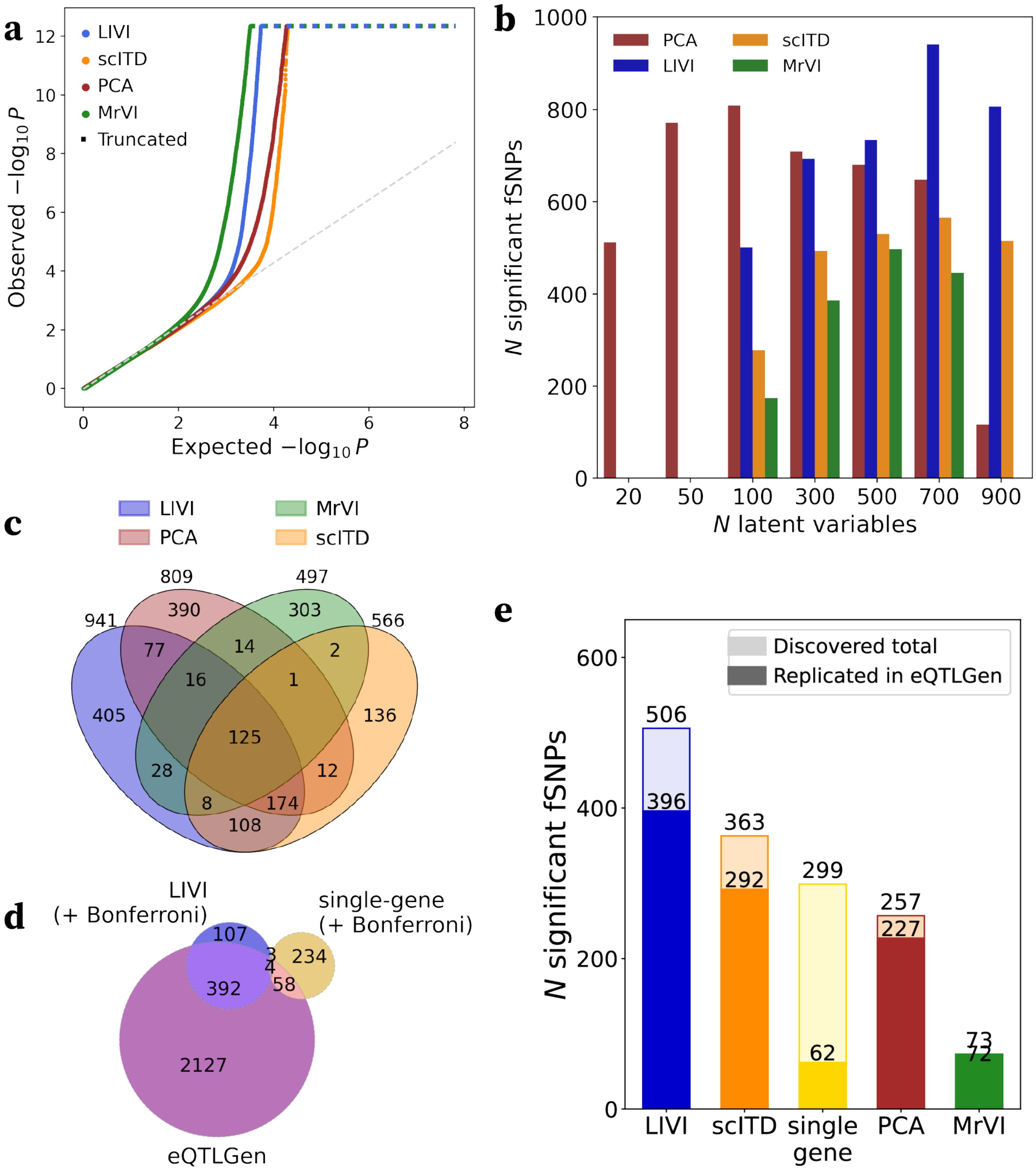

**Figure S11. Additional benchmarking results. (a)** Quantile-quantile (Q-Q) plot of the association $p$-values between SNPs and latent variables for the different latent variables methods: LIVI, scITD (Mitchel 2024), MrVI (Boyeau 2025) and PCA. For each model, the number of latent variables that maximized the number of discoveries was considered. **(b)** LIVI, scITD, MrVI and PCA were run with different numbers of latent factors/topics/PCs, and power to discover fSNPs was assessed in each setting. The y-axis shows the number of identified fSNPs, while the x-axis shows the number of latent variables tested for SNP effects. **(c)** Overlap between SNPs discovered by LIVI, scITD and MrVI and PCA. For each model, the number of latent variables that maximized the number of discoveries was considered. **(d)** Overlap between SNPs discovered by LIVI vs. gene-level testing after adjusting for the number of phenotypes tested with each approach. Additionally, the overlap with *trans*-eQTL variants identified in eQTLGen is shown (FDR $< 5\%$). **(e)** Number of variants in association with at least one latent variable (for LIVI, PCA, scITD, and MrVI) or gene after adjusting for the vast difference in the number of phenotypes (latent variables or genes) tested (Methods). Briefly, a two-step multiple testing correction procedure was employed, where we first applied Bonferroni correction for the number of phenotypes used in the association testing for each method and then applied Benjamini-Hochberg across SNP-phenotype pairs to maintain the FDR at 5%. For LIVI, PCA, scITD, and MrVI, the number of latent variables that maximized the number of discoveries was considered. Darker colors depict variants replicated in eQTLGen (Vosa 2021).

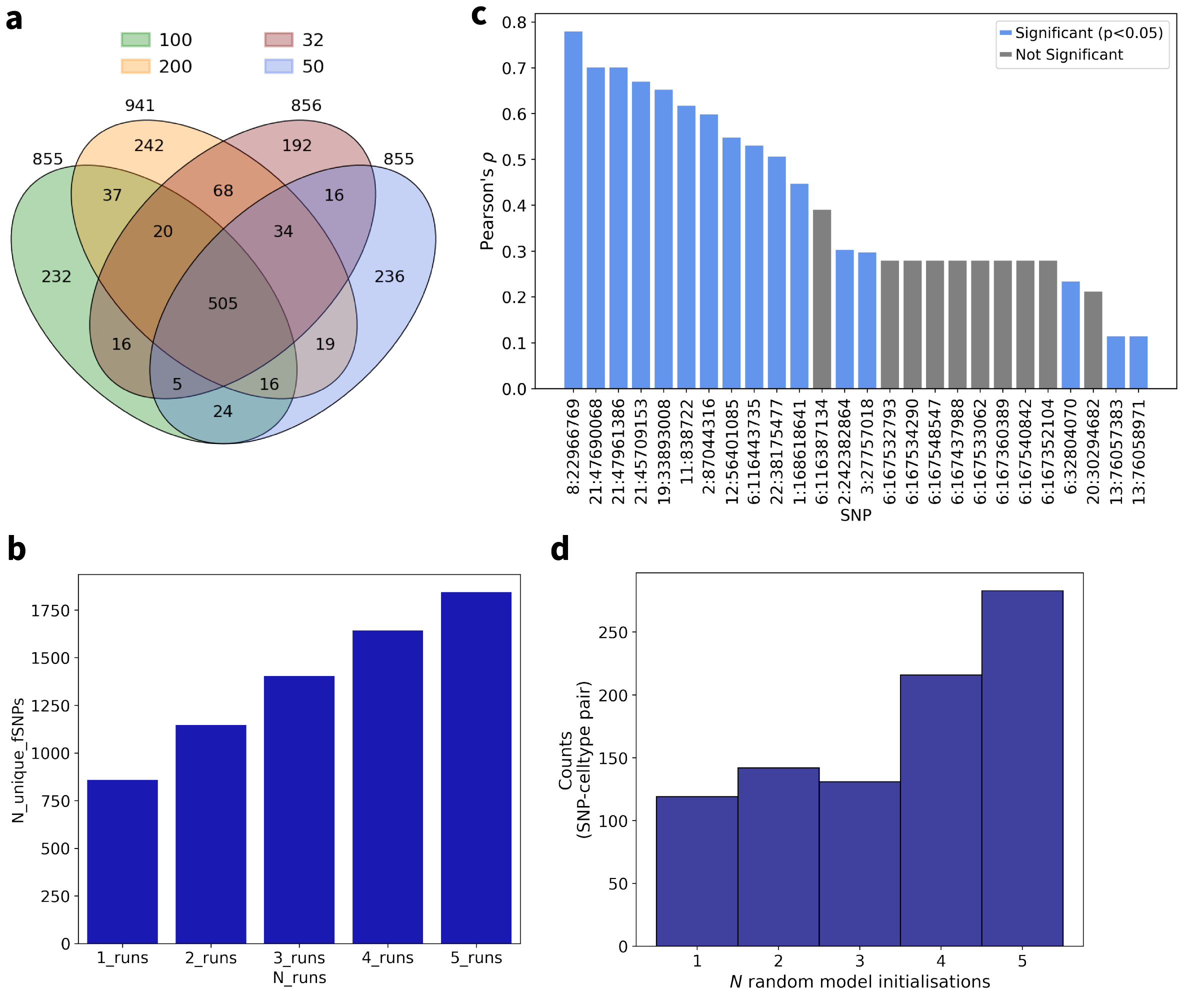

**Figure S12. fQTL robustness across model runs. (a)** Overlap of fQTLs discovered across different random restarts (seeds: 32, 50, 100, 200). **(b)** Aggregated number of unique fQTLs discovered by each additional model run. **(c)** Correlation of SNP effects at the single-cell level across different model runs (Methods). Shown are the mean correlations and $p$-values across all possible run pairs. **(d)** Concordance between fSNP to cell type assignments across different model runs. $D$ factors associated with the fSNP are assigned to one pre-annotated cell type as described in Methods.

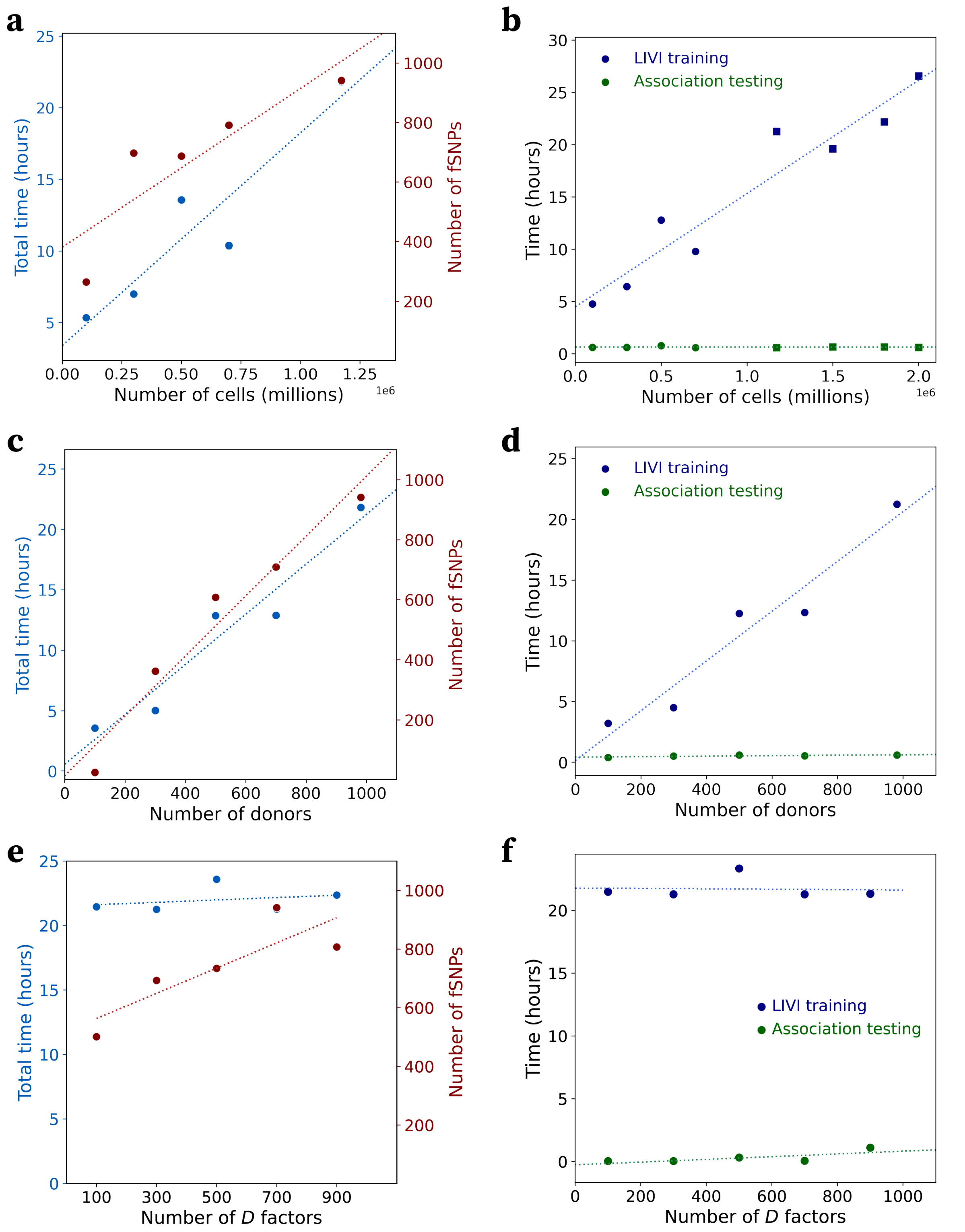

**Figure S13. Power and runtime as a function of the number of cells, donors and $D$ factors.** We assessed power and runtime across varying dataset sizes by sub- or oversampling the OneK1K (Yazar 2022) dataset. In all panels, the model architecture and number of cells or donors is the same as used in the main results unless otherwise indicated. **(a)** Runtime (blue) and number of significantly associated SNPs (red) for increasing number of cells. **(b)** Decomposition of total runtime to model training time (blue) and subsequent association testing time (green). Illustrated is the effect of the number of cells to each of the two time components. Squares denote models applied on artificially augmented datasets. **(c)** Runtime (blue) and number of significantly associated SNPs (red) for increasing number of donors, while keeping the number of $D$ factors fixed. Note that the number of cells is not fixed, as removing donors inherently removes cells as well. **(d)** Decomposition of total runtime to model training time (blue) and subsequent association testing time (green). Illustrated is the effect of the number of donors to each of the two time components. **(e)** Runtime (blue) and number of significantly associated SNPs (red) as a function of the number of $D$ factors used to train LIVI. **(f)** Decomposition of total runtime to model training time (blue) and subsequent association testing time (green). Illustrated is the effect of the number of $D$ factors to each of the two time components.

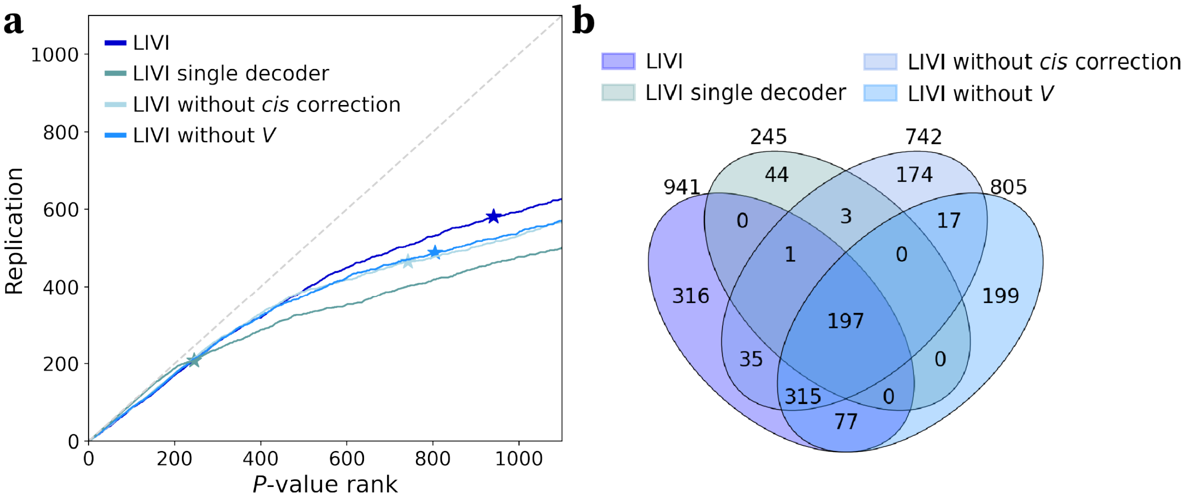

**Figure S14. Ablation studies. (a)** SNP $p$-value rank vs. replication in eQTLGen (Vosa 2021) for the full LIVI model (*LIVI*), as well as three reduced model formulations: one without global donor latent factors $\mathbf{V}$ (*LIVI without V*), one without correction for known *cis*-eQTLs (*LIVI without cis correction*), and one with a single decoder for cell-state and $D \times C$ factors (*LIVI single decoder*). Stars indicate the significance cut-off. **(b)** Venn diagram showing the overlap of discovered SNPs between the different variations of LIVI models described in **(a)**.

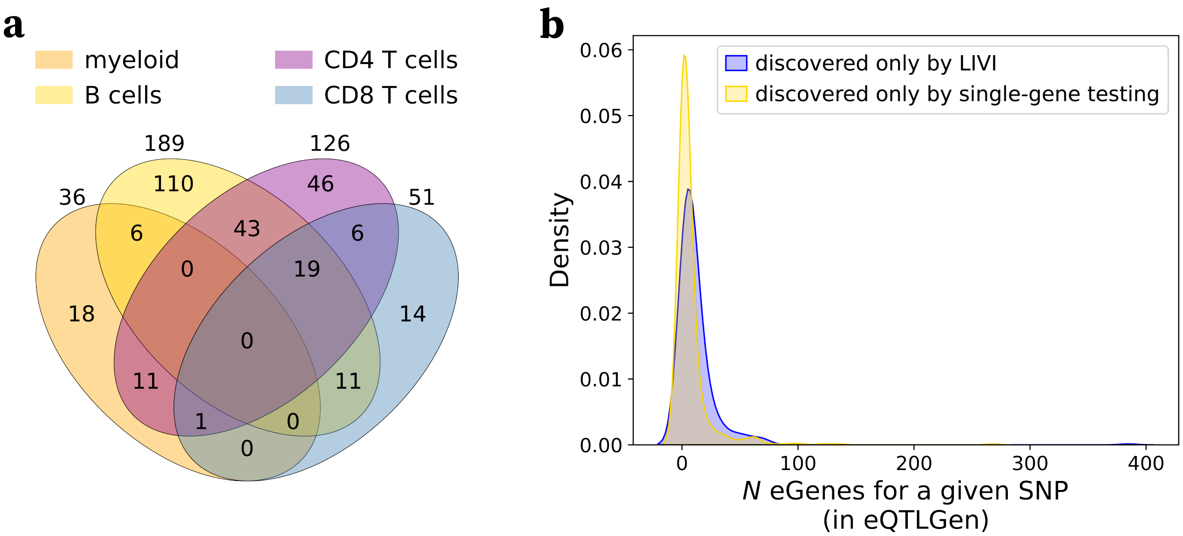

**Figure S15. fSNPs discovered by LIVI act in continuous cell states and/or affect the expression levels of a greater number of genes. (a)** Overlap between fSNPs estimated by LIVI to have an effect in myeloid cell differentiation trajectory(orange), naive B to plasma cell differentiation trajectory (yellow), naive to effector memory $CD4^+$ T cell differentiation trajectory (purple), or naive to effector memory $CD8^+$ T cell differentiation trajectory (blue). **(b)** Comparison of the number of corresponding eGenes for fSNPs identified exclusively by LIVI (blue) vs. eSNPs identified exclusively by gene-level testing (yellow), assessed using independent data (eQTLGen).

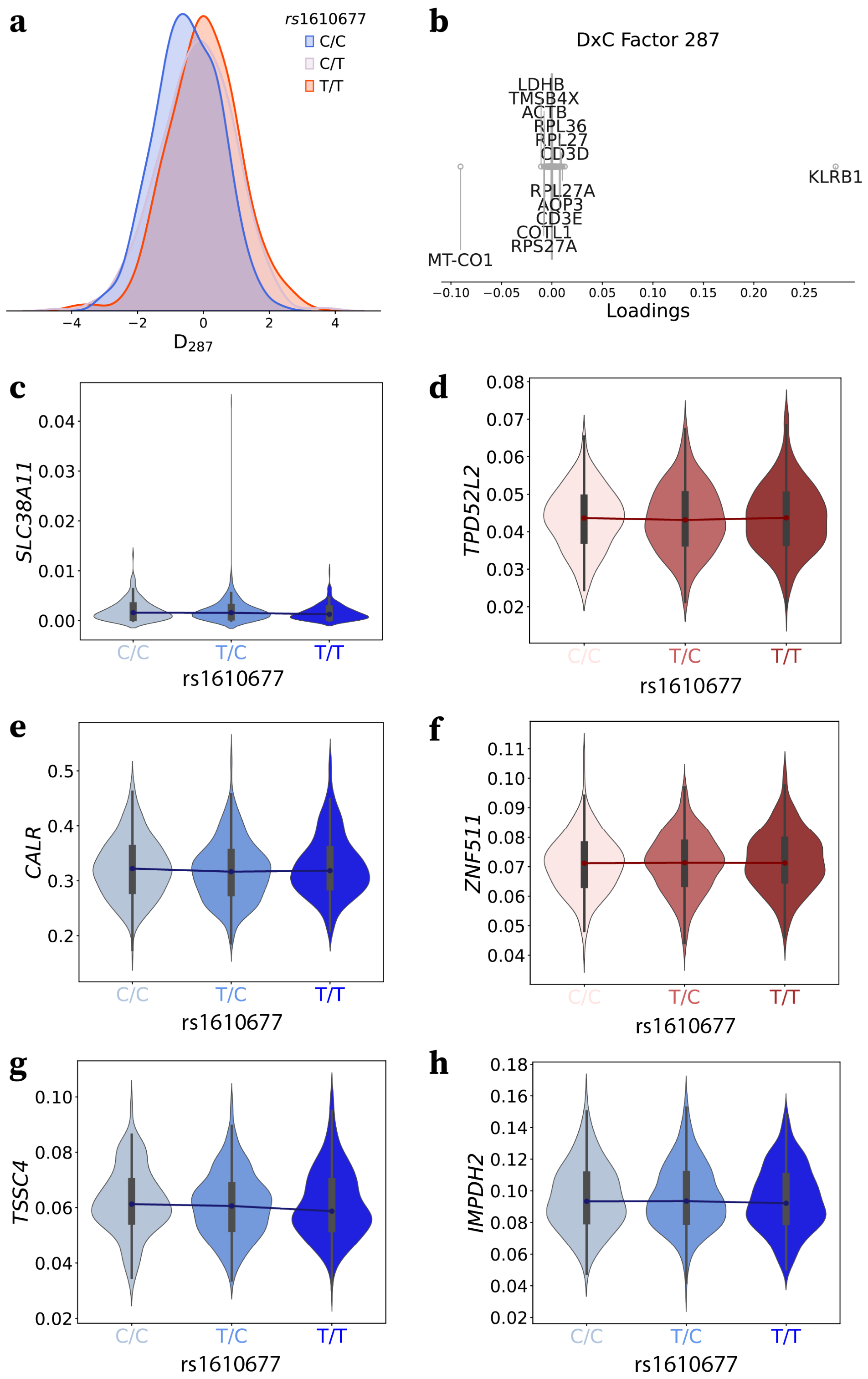

**Figure S16. LIVI discovers SNPs acting in cells differentiating from a CD4$^+$ T naive to an an effector memory phenotype.** **(a)** Distribution of $D_{287}$ values, stratified by donor genotype at rs1610677. **(b)** Boxplot of gene loadings for $D \times C_{287}$. **(c-h)** Distribution of the observed log-normalized expression of *trans* eGenes of rs1610677 discovered in eQTLGen (FDR $<$ 5%), stratified by donor genotype at rs1610677: **(c)** *SLC38A11*, **(d)** *TPD52L2*, **(e)** *CALR*, **(f)** *ZNF511*, **(g)** *TSSC4*, **(h)** *IMPDH2*. For each gene, donor-level expression was computed as the mean across all cells from that donor. Black boxes indicate the interquartile range (IQR), while whiskers mark the 1.5×IQR threshold.

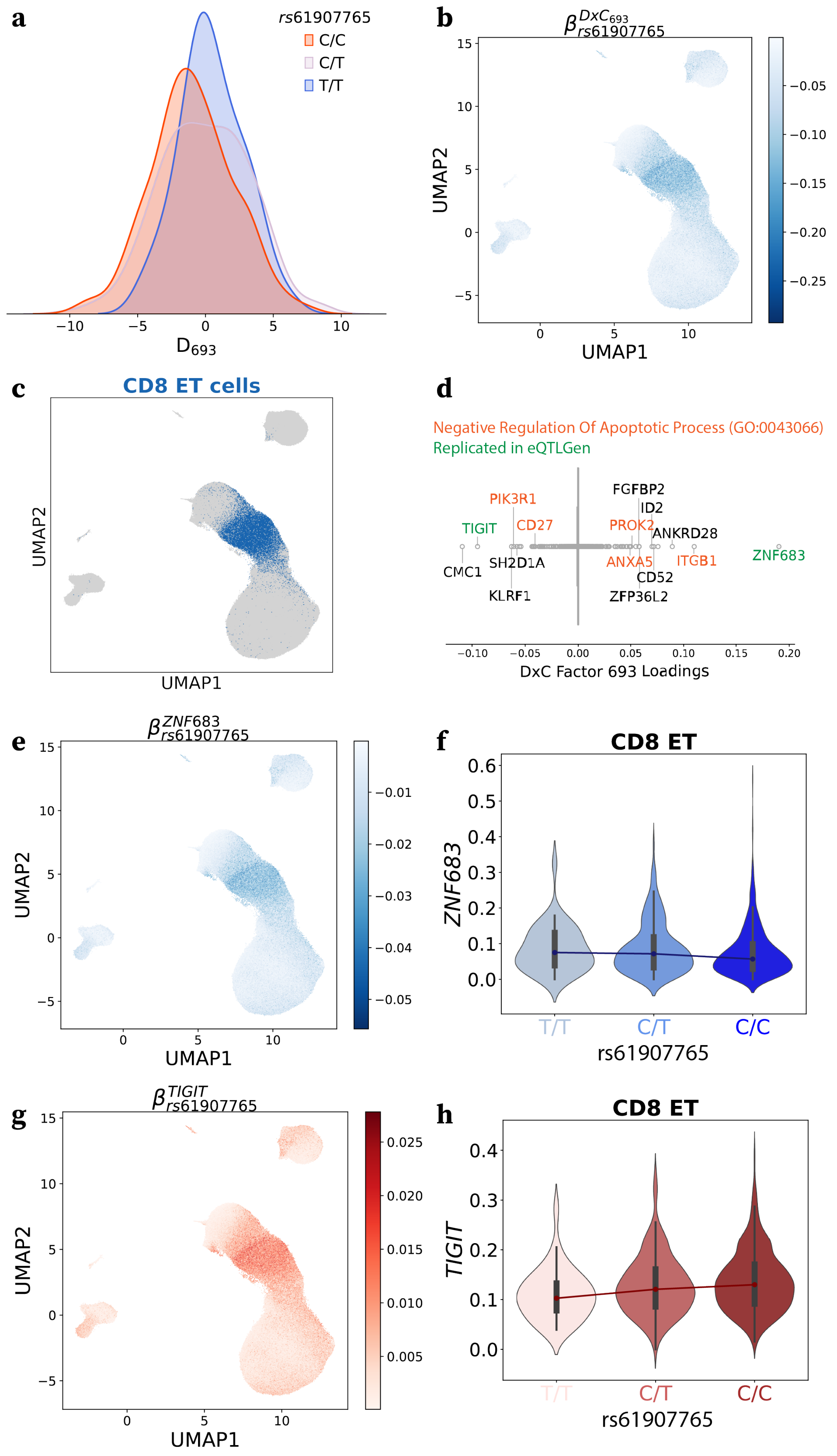

**Figure S17. LIVI groups genes known to be affected by the same SNP into the same factor.** **(a)** Distribution of $D_{693}$ values, stratified by donor genotype at rs61907765. **(b)** UMAP of LIVI's cell-state latent factors. Color denotes the estimated effect of rs61907765 on $D \times C_{693}$. **(c)** UMAP of LIVI's cell-state latent factors, highlighting cells annotated as effector memory CD8$^+$ T cells (CD8 ET) in the original publication (Yazar 2022). **(d)** Boxplot of gene loadings for $D \times C_{693}$. Genes involved in "Negative Regulation Of Apoptotic Process" (GO:0043066) are annotated in orange, while genes also found to be associated with rs61907765 in eQTLGen (Vosa 2021) in green. **(e,g)** UMAP on LIVI's cell-state latent factors. Color denotes the estimated effect of rs61907765 on the expression of **(e)** *ZNF683*, or **(g)** *TIGIT* at the single-cell level. **(f,h)** Distribution of the observed log-normalized expression of **(f)** *ZNF683*, or **(h)** *TIGIT* pseudobulked across CD8 ET cells and stratified by donor genotype at rs61907765. Black boxes indicate the interquartile range (IQR), while whiskers mark the $1.5\times$IQR threshold.

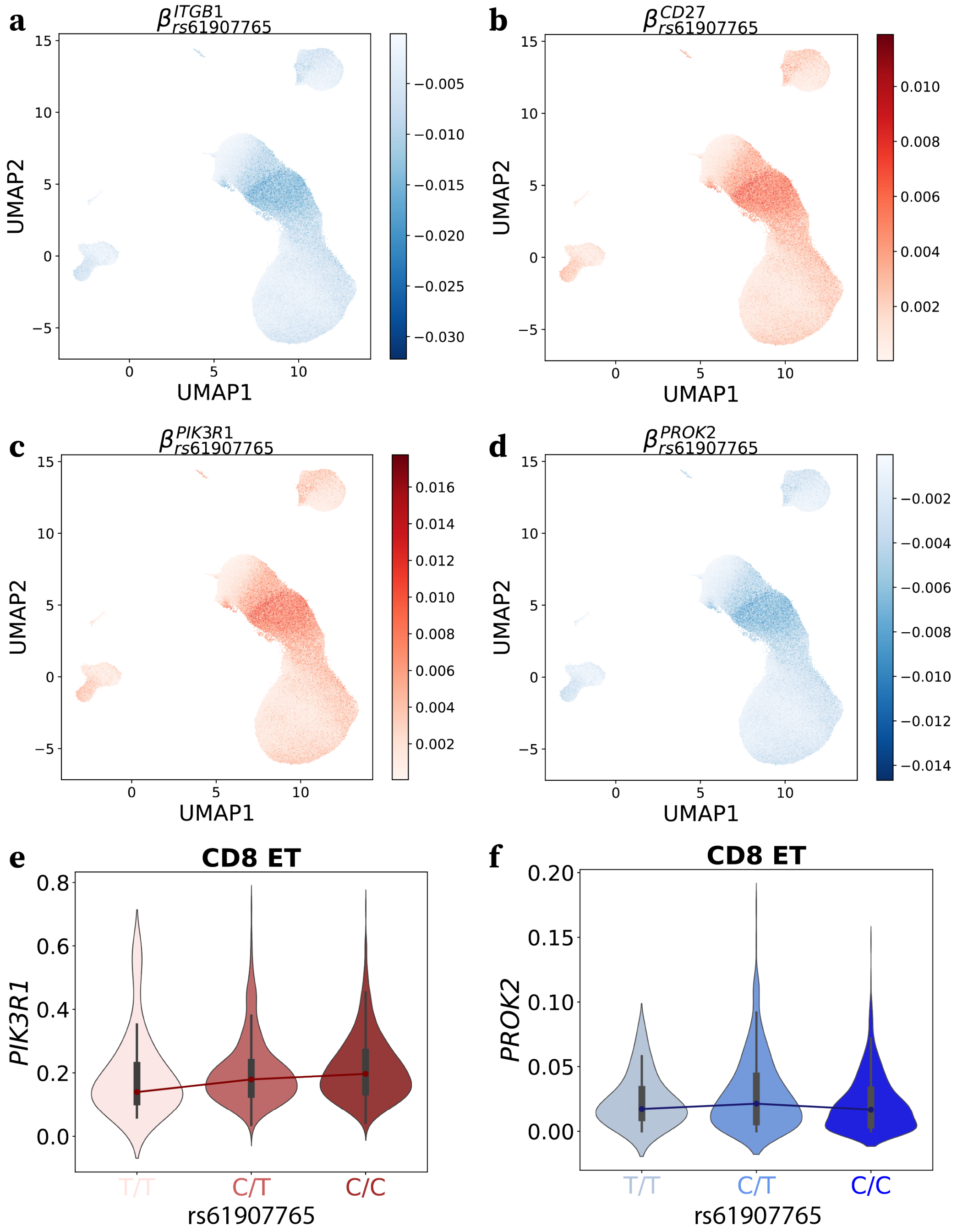

**Figure S18. LIVI identifies SNPs associated with expression of genes within the same biological process. (a-d)** UMAP on LIVI's cell-state latent factors. Color denotes the estimated effect of rs61907765 on the expression of **(a)** *ITGB1*, **(b)** *CD27*, **(c)** *PIK3R1*, or **(d)** *PROK2* at the single-cell level. **(e-f)** Distribution of the observed log-normalized expression of **(e)** *PIK3R1* or **(f)** *PROK2*, pseudobulked across CD8 ET cells and stratified by donor genotype at rs61907765. Black boxes indicate the interquartile range (IQR), while whiskers mark the 1.5×IQR threshold.

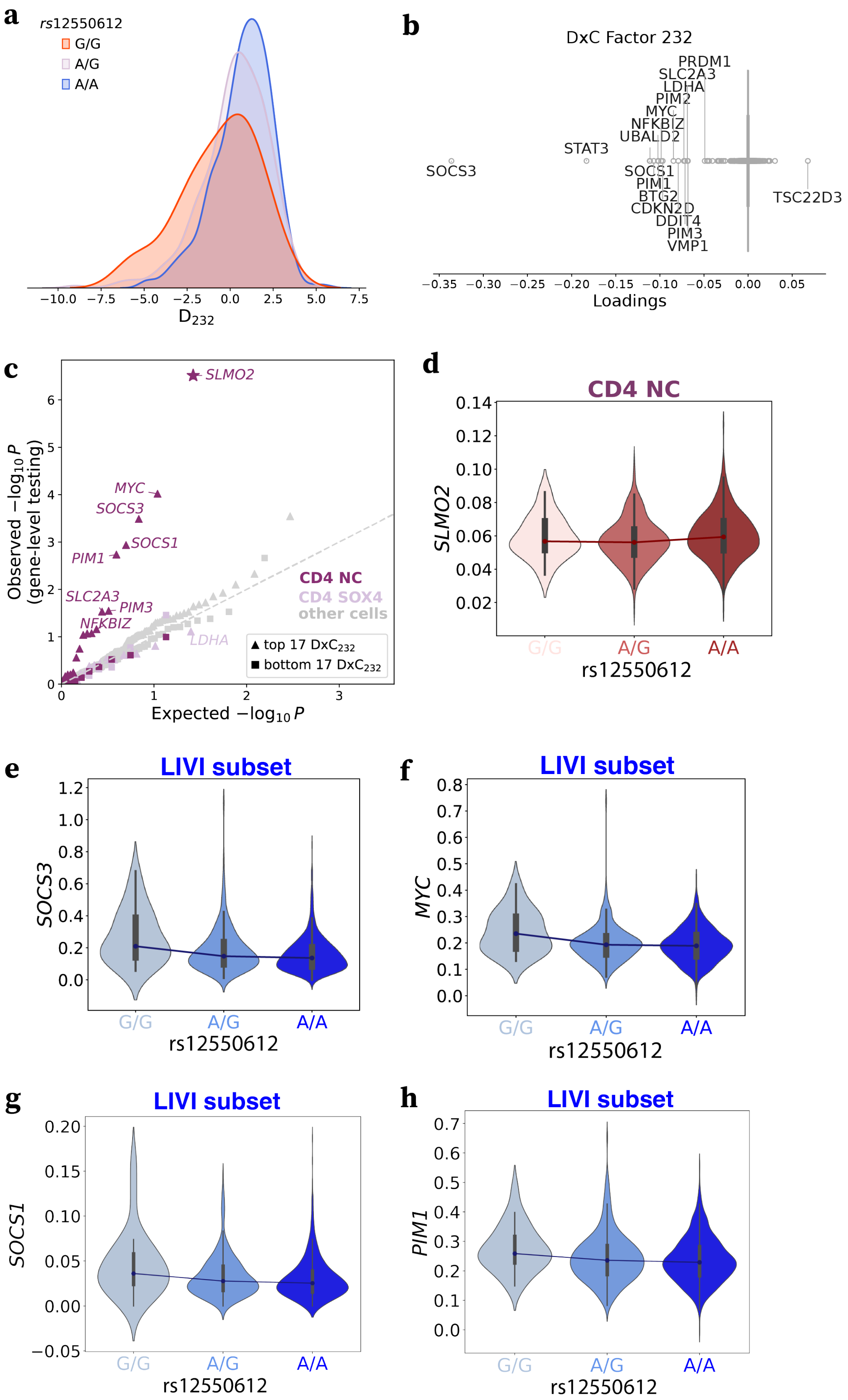

**Figure S19. LIVI uncovers gene-program-level effects of known eSNPs.** **(a)** Distribution of $D_{232}$ values, stratified by donor genotype at rs12550612. **(b)** Boxplot of gene loadings for $D \times C_{232}$. **(c)** Q-Q plot of expected vs. observed *p*-values for association tests between rs12550612 and the 17 genes with highest (triangles), as well as lowest (squares) absolute loadings for D $\times$ $C_{232}$, evaluated using single-gene tests in different cell types. Additionally, *SLMO2/PRELID3B*, the only gene discovered by single-gene tests to be significantly associated with rs12550612 (FDR $<$ 5%), is also included (star). Color indicates cell type (purple: CD4 NC, light purple: CD4 SOX4, grey: all other cells). **(d-h)** Distribution of the observed log-normalized expression, stratified by donor genotype at rs12550612. Black boxes indicate the interquartile range (IQR), while whiskers mark the 1.5$\times$IQR threshold. (d) Pseudobulked *SLMO2* expression across CD4 NC cells for each donor. (e-h) Pseudobulked (e) *SOCS3*, (f) *MYC*, (g) *SOCS1*, (h) *PIM1* expression across *LIVI subset* cells for each donor.

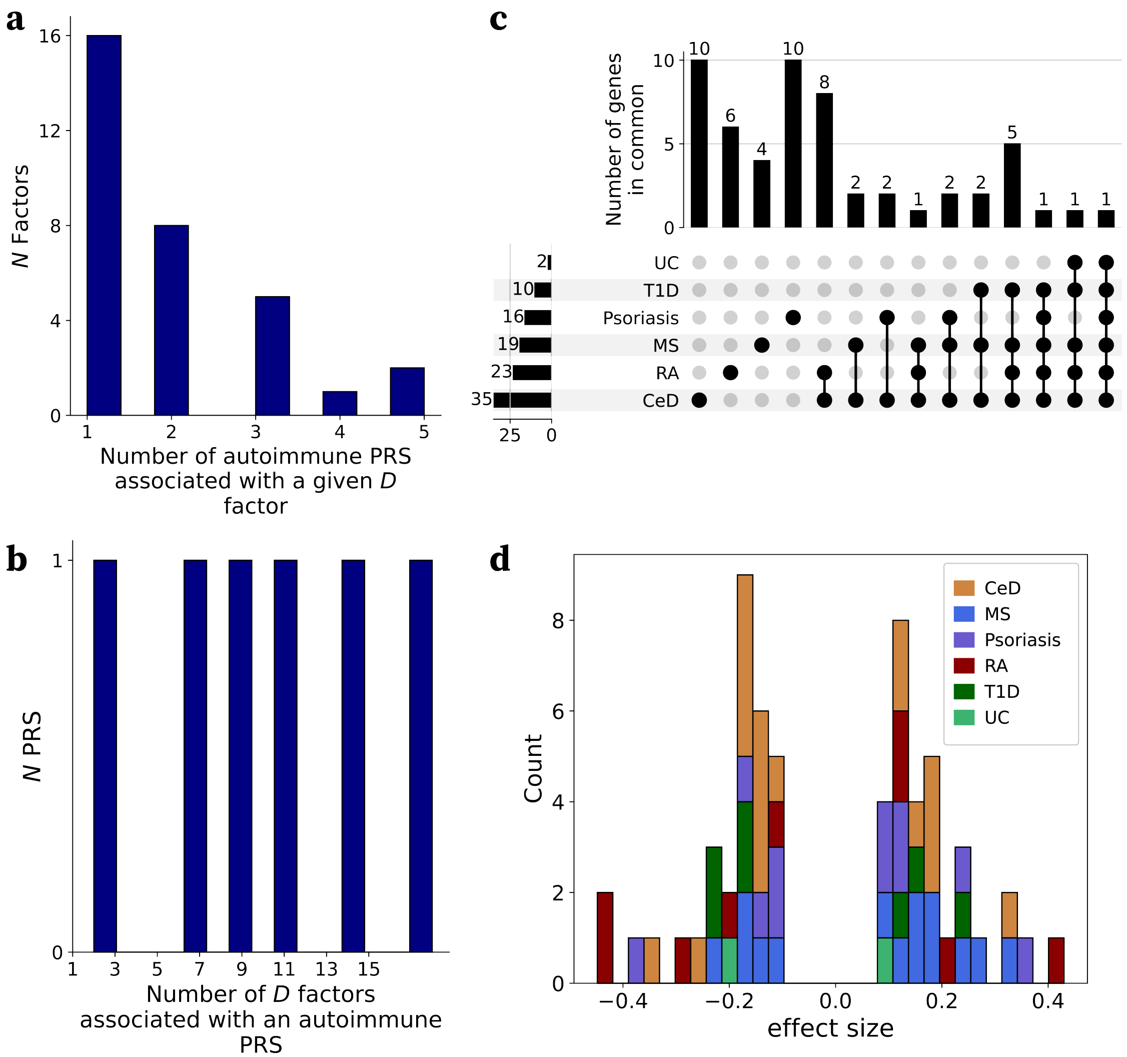

**Figure S20. PRS for autoimmune diseases are associated to multiple $D$ factors. (a)** Number of unique autoimmune PRS a given $D$ factor is associated with. **(b)** Number of unique $D$ factors a given autoimmune PRS is associated with. **(c)** Upset plot of genes estimated by LIVI to be involved in the polygenic risk for autoimmune diseases. **(d)** Histogram showing the distribution of effect sizes for PRS–factor associations across autoimmune diseases. Each color represents a different disease: celiac disease (CeD), multiple sclerosis (MS), psoriasis, rheumatoid arthritis (RA), type 1 diabetes (T1D), and ulcerative colitis (UC).

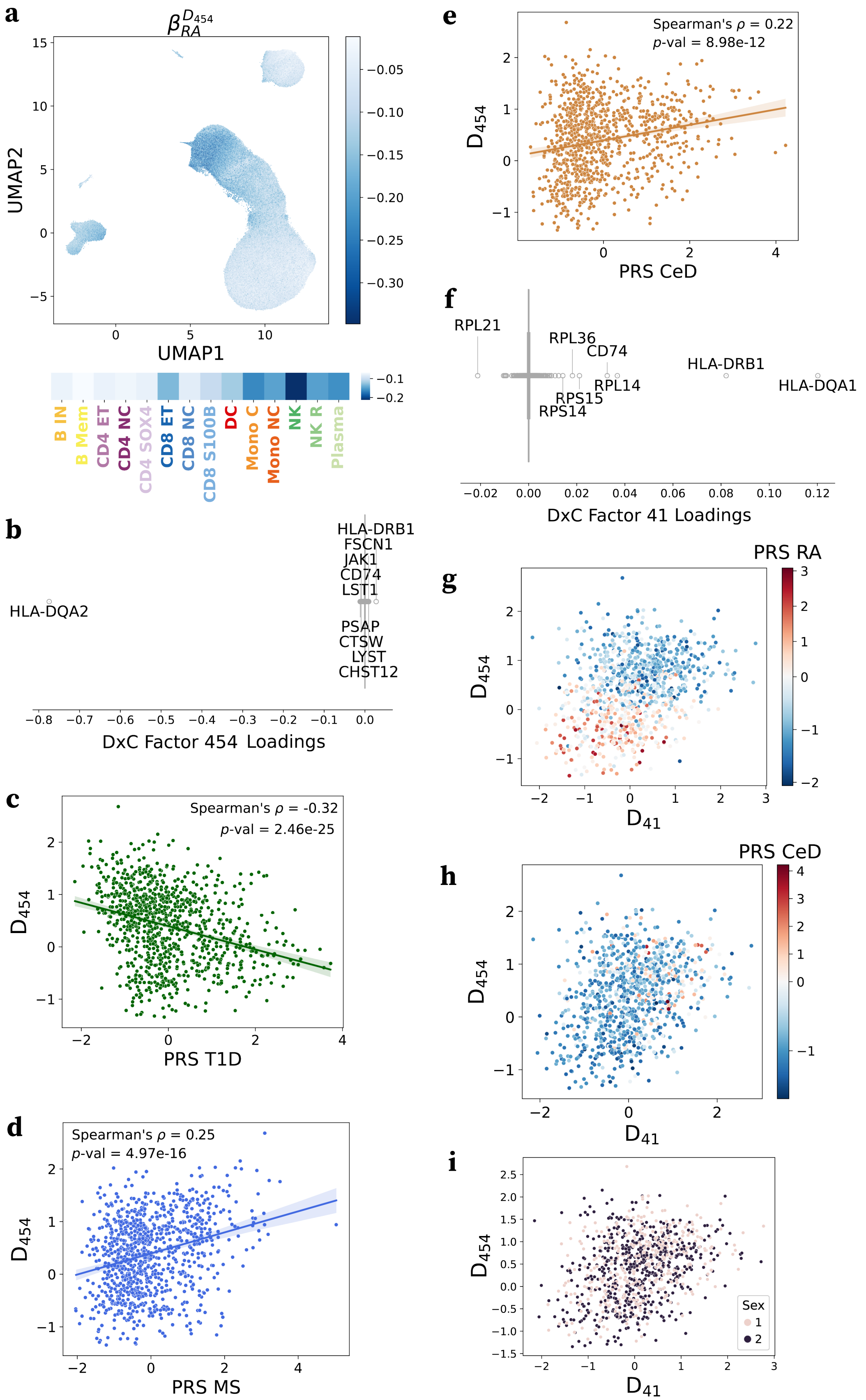

**Figure S21. LIVI donor factors capture polygenic disease risk. (a)** Top: UMAP of LIVI's cell-state latent factors. Color denotes the estimated effect of PRS for rheumatoid arthritis (RA) on $G \times C_{454}$. Bottom: Estimated effect of PRS for RA on $G \times C_{454}$ averaged across cells of different types from the original publication (Yazar 2022). **(b)** Boxplot of gene loadings for $D \times C_{454}$. **(c-e)** Scatterplot between scaled PRS for **(c)** type 1 diabetes (T1D), **(d)** multiple sclerosis (MS), or **(e)** celiac disease (CeD) vs. $D_{454}$ activity in each donor. Spearman's $\rho$ between PRS for (c) T1D, (d) MS, or (e) CeD and $D_{454}$ and corresponding $p$-value are noted. **(f)** Boxplot of gene loadings for $D \times C_{41}$. **(g-i)** Scatterplot between $D_{41}$ vs. $D_{454}$. Color denotes **(g)** scaled PRS for RA, **(h)** scaled PRS for CeD, or **(i)** the donor sex.
